# Distinct *Escherichia coli* transcriptional profiles in the guts of recurrent UTI sufferers revealed by pangenome hybrid selection

**DOI:** 10.1101/2024.02.29.582780

**Authors:** Mark G. Young, Timothy J. Straub, Colin J. Worby, Hayden C. Metsky, Andreas Gnirke, Ryan A. Bronson, Lucas R. van Dijk, Christopher A. Desjardins, Christian Matranga, James Qu, Jesús Bazan Villicana, Philippe Azimzadeh, Andrew Kau, Karen W. Dodson, Henry L. Schreiber, Abigail L. Manson, Scott J. Hultgren, Ashlee M. Earl

**Author notes:** These authors contributed equally to this work (Mark G. Young, Timothy J. Straub).

## Abstract

Low-abundance members of microbial communities are difficult to study in their native habitats. This includes *Escherichia coli*, a minor, but common inhabitant of the gastrointestinal tract and opportunistic pathogen, including of the urinary tract, where it is the primary pathogen. While multi-omic analyses have detailed critical interactions between uropathogenic *Escherichia coli* (UPEC) and the bladder that mediate UTI outcome, comparatively little is known about UPEC in its pre-infection reservoir, partly due to its low abundance there (<1% relative abundance). To accurately and sensitively explore the genomes and transcriptomes of diverse *E. coli* in gastrointestinal communities, we developed *E. coli* PanSelect which uses a set of probes designed to specifically recognize and capture *E. coli*’s broad pangenome from sequencing libraries. We demonstrated the ability of *E. coli* PanSelect to enrich, by orders of magnitude, sequencing data from diverse *E. coli* using a mock community and a set of human stool samples collected as part of a cohort study investigating drivers of recurrent urinary tract infections (rUTI). Comparisons of genomes and transcriptomes between *E. coli* residing in the gastrointestinal tracts of women with and without a history of rUTI suggest that rUTI gut *E. coli* are responding to increased levels of oxygen and nitrate, suggestive of mucosal inflammation, which may have implications for recurrent disease. *E. coli* PanSelect is well suited for investigations of native *in vivo* biology of *E. coli* in other environments where it is at low relative abundance, and the framework described here has broad applicability to other highly diverse, low abundance organisms.

## Introduction

Urinary tract infections (UTIs) are among the most common bacterial infections worldwide, with significant individual and societal impacts. Recurrent UTIs (rUTIs) affect millions, mostly women ^1^, and drain precious antibiotic resources ^2–6^. The vast majority of UTIs are caused by uropathogenic *Escherichia coli* (UPEC)^7^. In contrast to other *E. coli* pathotypes, such as enterohemorrhagic *E. coli* (EHEC), the genetic definition of UPEC has remained elusive, given the wide diversity of *E. coli* that cause UTIs and rUTIs ^8–10^. *E. coli* are prevalent but low-abundance (<1% relative abundance) members of the gastrointestinal tract (gut) from where they can be excreted in feces and go on to colonize the bladder ^11–13^. Culture and metagenomic sequencing-based investigations have revealed much about this functionally and evolutionarily diverse group, including the carriage of multiple *E. coli* strains in the human gut at once ^14^. However, these approaches have drawbacks that leave open key questions about UPEC lifestyle, including the interplay between UPEC in the gut and infection of the bladder.

Advances in selective nucleic acid enrichment prior to sequencing have made it possible to study low abundance organisms in their native habitats at high resolution. Solution hybrid selection (HS) ^15^, also called hybrid capture, relies on biotinylated oligonucleotides, or probes, that selectively hybridize (*i.e.*, bind) to target sequences, effectively tagging them for capture on streptavidin beads. Originally developed for human exome sequencing ^16^, HS has also enabled sequencing malarial ^17,18^ and viral ^19^ genomes despite an excess of human host material in the sample. Since HS preserves the relative abundance of cDNAs in RNA-Seq libraries^20^, it has also been used for transcriptomic applications, such as expression profiling of *Bacteroides fragilis* from murine intestinal compartments^21^. While these prior studies have focused on individual target organisms, or species with limited overall diversity, many bacterial species, like *E. coli*, have large pangenomes (i.e., the entire set of genes found across a species) that cannot be represented by a single reference genome. Having emerged >100 million years ago from a common ancestor of *Salmonella* ^22^, *E. coli* have evolved considerably with more than eight recognized distinct phylogroups^23^, collectively accumulating a pangenome estimated at over 120,000 gene families ^24^. This functional variation can have broad implications for a strain’s physiology, interactions, and disease potential^14^.

To increase our ability to understand UPEC in their native gut habitat, and in the context of rUTI, we developed an HS probe set for enrichment of the *E. coli* pangenome, using a combination of comparative genomics and a recently developed algorithm^25^ that enables HS probe design for efficient tiling across diverse sequences (**Figure 1)**. Our “*E. coli* PanSelect” method selectively enriches, by orders of magnitude, diverse *E. coli* genomes and transcriptomes present at low abundance in complex communities. Applied to human stool from a clinical cohort study of rUTI, *E. coli* PanSelect revealed some of the adaptive strategies deployed by *E. coli* while residing in the rUTI gut, including a shift from fermentative to aerobic metabolism, which may have implications for understanding and treating recurrent disease. The pangenome-based HS approach described here holds promise for investigating other diverse taxa in a wide variety of community contexts.

**Figure 1.**
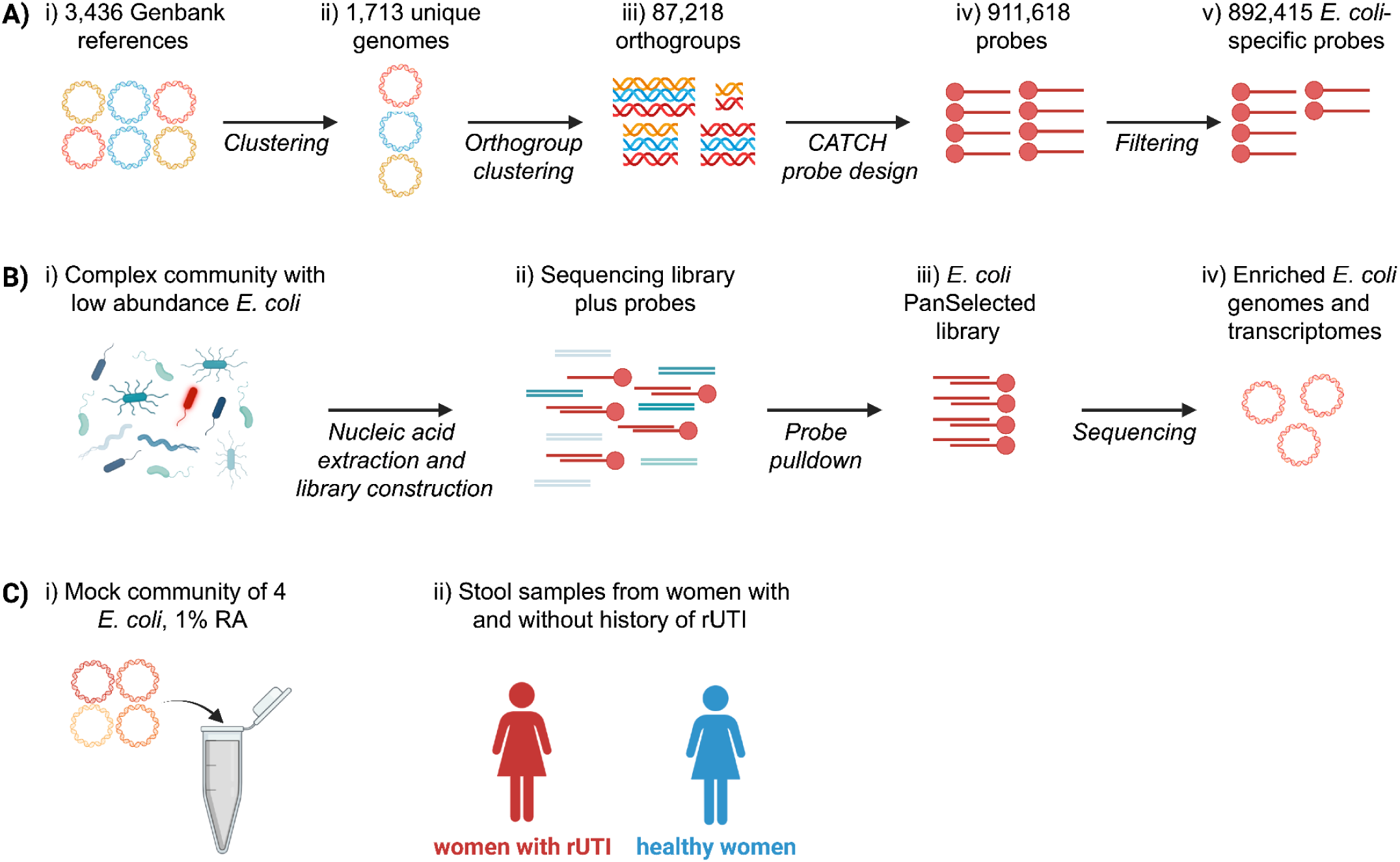
*E. coli* PanSelect probe design and applications. a) Probe design. i) All available, complete *E. coli* genomes were downloaded from RefSeq (295) and the NCBI Pathogens database (3,141). ii) k-mer similarity was used to identify 1,713 unique genome clusters. iii) Orthologous gene groups were constructed from these genome clusters with SynerClust ^71^, filtered based on prevalence, and further clustered at 80% identity with UCLUST ^72^. 60-75 bp probes with specificity to the resulting clusters were iv) generated with CATCH^25^ and v) filtered based on homology to other common gut microbes (ie *Bacteroidetes* and *Firmicutes*). b) *E. coli* PanSelect workflow. i) Sequencing libraries are constructed from complex communities containing low abundances of *E. coli* (red). ii) Short, biotinylated oligonucleotide probes are added to the sequencing library, which bind complementary sequences. iii) Streptavidin pulldown is used to isolate bound target sequences from the library before iv) sequencing. c) Applications of *E. coli* PanSelect. i) Enrichment of a four strain mock community for initial benchmarking. ii) Analysis of *E. coli* gene content and transcription in stool from a clinical study of recurrent UTIs (rUTI). *Created with BioRender.com*.

## Results

### Probe set design covers vast majority of *E. coli* pangenome

In order to design an HS probe set to cover known *E. coli* genetic diversity, we downloaded all of the available 3,436 high-quality *E. coli* assemblies from NCBI’s RefSeq and Genbank databases and then clustered them to a set of 1,713 unique *E. coli* references spanning all major phylogroups (**Figure 1**, **Supplementary** Figure 1**, Supplementary Table 1-2**; **Methods**). We used the CATCH algorithm for HS probe design^25^. To reduce computational complexity, we provided CATCH with refined orthologous gene clusters (80% or higher nucleotide identity) containing three or more genes or a Pfam domain of interest (**Supplementary table 13; Methods**). CATCH designed 911,618 probes of length 60-75bp, which we filtered to a final set of 892,415, based upon predicted off-target specificity to other common gut bacteria (e.g., *Bacteroidetes*) (**Methods**).

We next performed an *in silico* experiment to determine how well the CATCH-designed probe set covered the *E. coli* pangenome by querying each probe sequence against all 8,334,026 genes from the full set of unique references. We classified a gene as ‘selected’ by a probe if it had a blast hit of at least 65 bp, and no more than 8 mismatches (**Methods**). A total of 97.92% of all genes in the *E. coli* pangenome were successfully selected, including 29,232 of the *E. coli* genes that were not included in the probe design (likely due to sequence homology). Of the 8,168,837 genes used in probe design, 99.95% were selected, with most non-selected genes having “hypothetical protein annotations.

### *E. coli* PanSelect enriched four strains of *E. coli* in a mock metagenomic community without biasing strain composition

As a first assessment of *E. coli* PanSelect, we enriched *E. coli* from an Illumina sequencing library created from a previously described mock community, containing an uneven mixture of DNA from four sequenced *E. coli* isolates in a background of human DNA ^26^. The sequence quality of pre- and post-HS datasets were comparable though the duplication rates were higher post HS, which we adjusted for prior to comparisons **(Supplementary Results A; Methods)**. *E. coli* PanSelect enriched the total relative abundance of *E. coli* 40-fold, without changing the strain composition of the community (paired t-test of RA ratios, p > 0.1, **Figure 2a, Methods**). The depth of *E. coli* genome coverage increased from 6x to 200x for the most abundant strain, and from 0.6x to 27x for the least abundant strain (**Figure 2b**). After enrichment, 72% (up from 0.8%) of the least abundant strain’s reference was covered with 5 or more reads (**Figure 2c)**. As expected, we were able to produce substantially more complete assemblies using the HS-enriched data (**Supplementary Table 4-5, Supplementary Results B**). We observed enrichment in regions up to ∼300 bp away from predicted probe hybridization sites (**Figure 2d**), leading to 97-99% of each strain’s genome being enriched by HS instead of the 83-86% which would be predicted based on probe coverage alone (**Table 1**).

**Figure 2.**
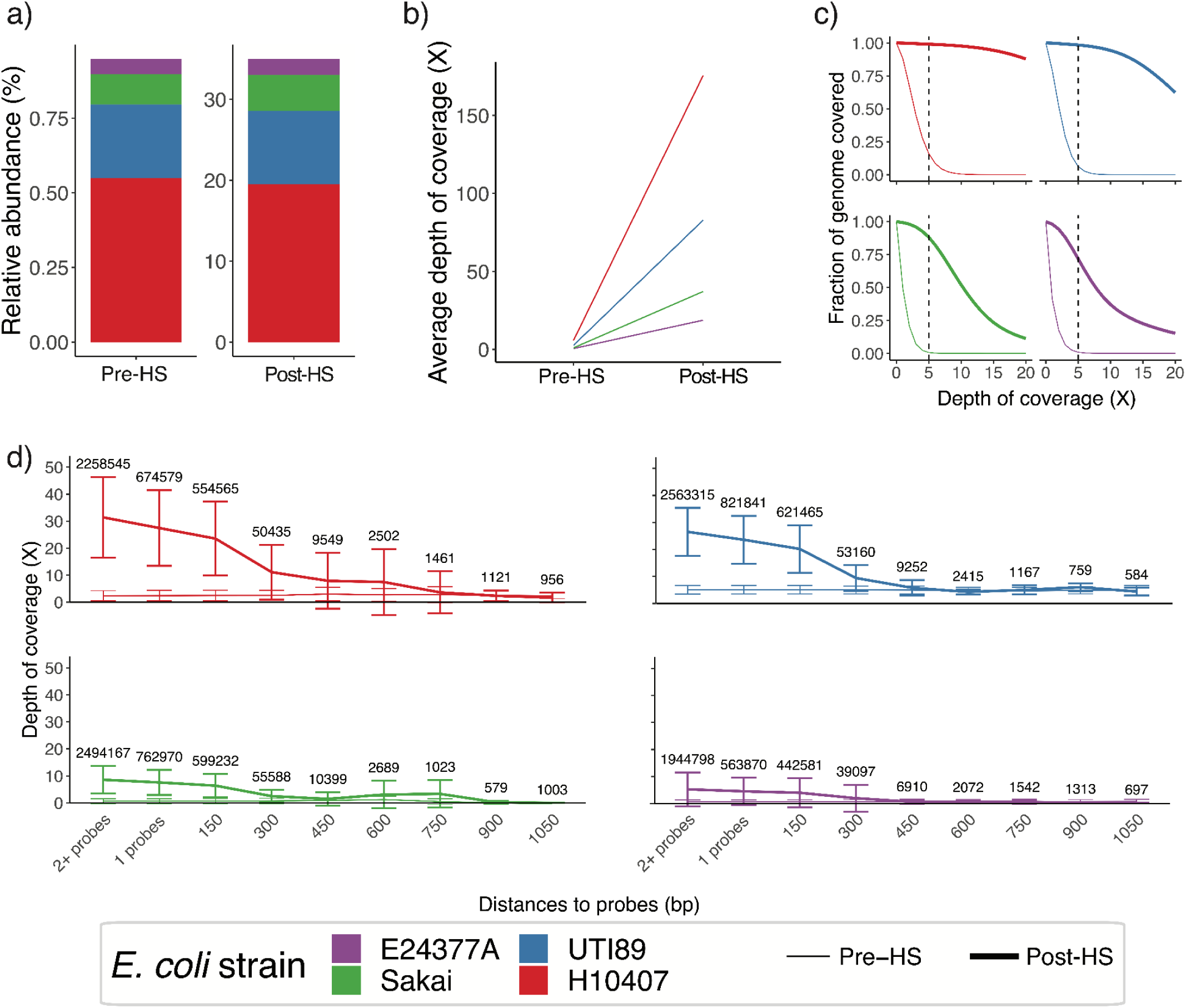
*E. coli* PanSelect enriches *E. coli* DNA without bias from a 4-strain mock community. a) The RAs of the four *E. coli* strains, pre- and post-HS, calculated using StrainGE. b) Average depth of coverage pre- and post-HS for each strain. c) Genome coverage pre-HS (thin lines) and post-HS (thick lines) for each strain. The dashed vertical line represents 5x coverage. d) Average depth of coverage in relation to the closest predicted probe binding site(s), for each of the four strains. “1 probe” and “2+ probes” indicate regions where probes are predicted to bind. Error bars denote standard deviations. Numbers above error bars indicate the number of positions across the genome in each category. Thin lines represent pre-HS data; thick lines post-HS data.

**Table 1.** Enrichment of strain genomes for the mock community: Post-HS depth of coverage for the mock community was increased for >96% of each strains’ genome.

|  | Depth of coverage |  |  |
| --- | --- | --- | --- |
| Strain | Unenriched |  | Enriched |
|  | Missing in both pre-HS and HS* | HS <= pre-HS | HS > pre-HS |
| H10407 | 0.02% | 0.58% | 99.40% |
| UTI89 | 0.05% | 0.70% | 99.25% |
| Sakai | 0.37% | 1.18% | 98.44% |
| E24377A | 1.51% | 1.62% | 96.86% |

\*Missing indicates neither pre-HS nor HS data had coverage. Other columns indicate HS data compared to pre-HS data depth of coverage.

### *E. coli* PanSelect provides highly enriched views of *E. coli* DNA and RNA within human stool samples without biasing the unenriched, background community

We next used *E. coli* PanSelect to enrich *E. coli* from 188 DNA-based and 130 RNA-Seq (cDNA-based) Illumina libraries (**Supplementary Results A, Supplementary Table 6),** constructed from human stool collected as part of a previously published clinical study investigating the gut as a uropathogen reservoir in recurrent urinary tract infection (rUTI) ^27^. Post-HS DNA libraries had a median 158-fold increase in *E. coli* RA (range of 5-2,232-fold increase) compared to pre-HS libraries (average of 0.07% *E. coli* RA), enabling the detection of one additional strain for every 2.8 samples (**Figure 3a**). *E. coli* genome assemblies from post-HS samples were vastly improved, including for one sample with a corresponding near-complete assembly of a cultured isolate. The amount of the genome that we were able to assemble from metagenomic reads from this sample increased from <1% pre-HS to >99% post-HS (**Supplementary Results B**, **Supplementary Tables 8-9**). As with the mock community, RA ratios between strains were not significantly biased in samples harboring multiple strains (**Figure 3b**; Wilcoxon signed-rank test, p=0.29 for subset of samples with two strains before and after enrichment).

**Figure 3.**
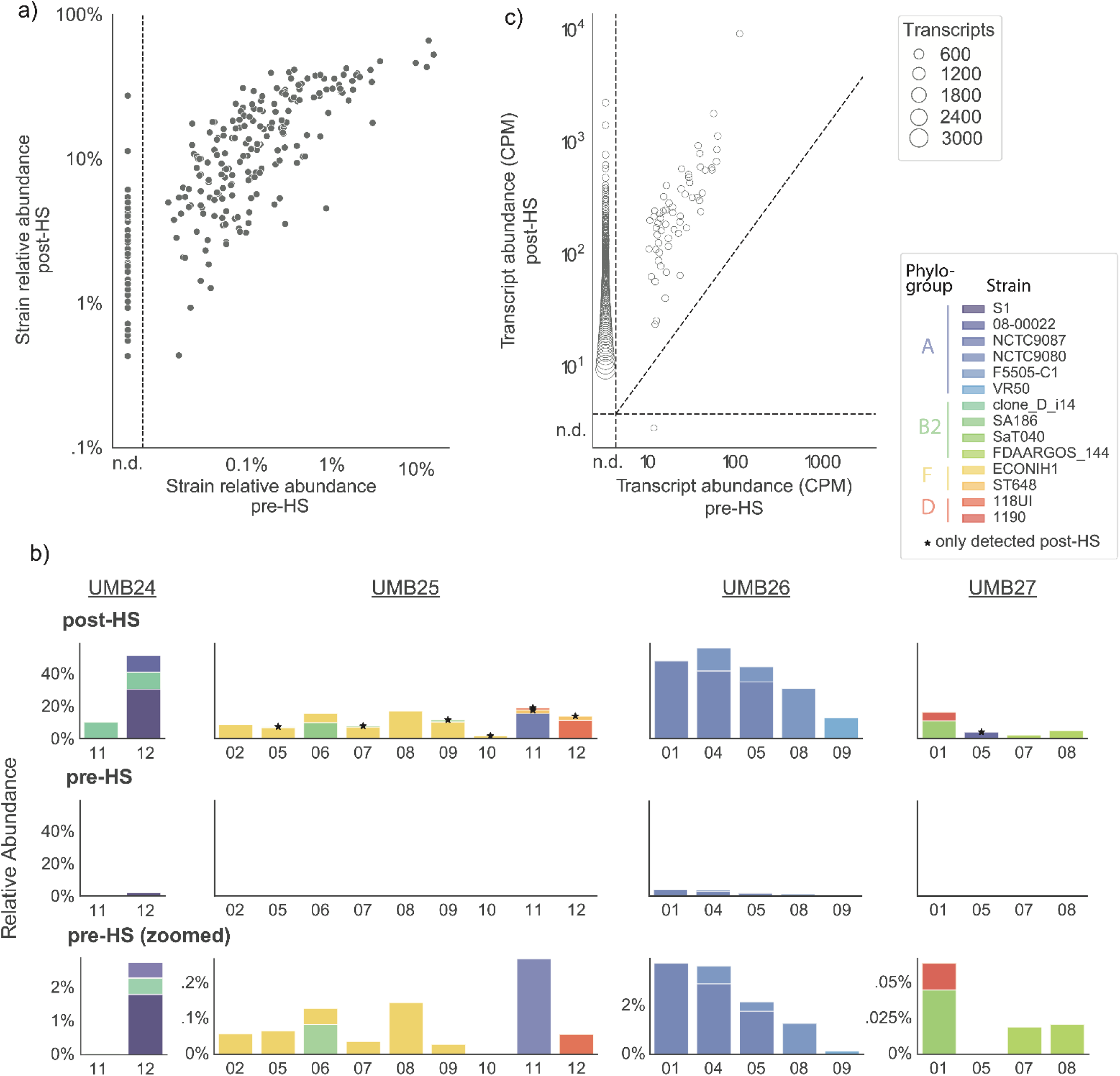
*E. coli* PanSelect enriched *E. coli* from human stool samples, revealing previously missed strains and transcripts. a) Pre- and post-enrichment RAs of *E. coli* strains detected within 188 human stool metagenomes. Points to the left of the dashed vertical line represent strains that were not detected (n.d.) within unenriched metagenomes. Strains were identified with StrainGST (Methods). b) Observed pre- and post-HS expression levels of individual transcripts, for the set 94 pre- and post-HS metatranscriptomes with at least 1 million reads, downsampled to 1 million reads. Transcripts expressed below 10 copies per million (CPM) are classified as not detected (n.d.). Points represent clusters of transcripts expressed at similar levels, formed with hierarchical clustering. c) Strain composition of samples from four randomly chosen participants, after enrichment (top row) and before enrichment (bottom two rows). Pre-HS data is shown using the same y-axis scale as post-HS data, as well as zoomed in to see the strain composition. Strain RAs were estimated with StrainGST. Stars indicate strains that could not be detected before HS.

Results were largely similar for RNA-Seq libraries. There was a median 30-fold increase in *E. coli* RA (range of 1.2- to-5,084-fold increase), from a median starting RA of 0.12%. In a comparison of metatranscriptomes downsampled to one million reads (**Methods**), we found that an average of 20x (728 vs 37) more unique genes were detected post-HS than pre-HS. There were only 41 instances of a transcript observed in a pre-HS sample that was not detected post-HS, 25 of which came from a single sample (UMB12_02) with a high pre-HS RA of *E. coli* (**Figure 3c**).

Post-HS libraries contained sequences from other microbes besides *E. coli* (83% for DNA- and 92% for RNA-Seq libraries), consistent with our prior experience using HS ^21^. To determine the utility of post-HS data to explore non-*E. coli* microbial fractions, we constructed taxa (DNA) and gene family (RNA) RA profiles with MetaPhlAn3 and HUMAnN3, respectively, and compared profiles between pre- and post-HS libraries. We observed only minor differences in the non*-E. coli* composition of the taxa and gene profiles between pre- vs. post-HS libraries (p> 0.05; PERMANOVA on Bray-Curtis dissimilarities), suggesting that *E. coli* PanSelect libraries could be used for analysis of the full microbiome in addition to the *E. coli* population (**Supplementary** Figure 2-3).

### *E. coli* gene content did not differ significantly between healthy women and those with rUTI

Though our prior work using unenriched stool metagenomes suggested that *E. coli* in healthy and rUTI guts are similar in their RAs and *E. coli* phylogroup membership ^27^, the sequencing coverage of *E. coli* genomes in these analyses was relatively scant. To explore this question with greater sensitivity, we used the *E. coli* PanSelect data described above to directly compare *E. coli* gene content between women with and without history of rUTI. Of the 13 participants in the rUTI cohort with measurable *E. coli* in at least one sample, we excluded four individuals who did not report UTI symptoms during the one-year study period to focus on patients with active disease (i.e., recurrers) (**Supplementary Table 7)**. Using PanPhlAn 3 to profile *E. coli* coding sequencing diversity at the level of UniRef90 clusters ^28^, we identified an average of 4,618 unique *E. coli* gene families per sample, roughly equivalent to the number of genes in a typical *E. coli* genome^29^. Individually, none of the ∼16,000 gene families detected across samples were significantly different in their occurrence in recurrer vs healthy cohort samples (**Supplementary Table 10; Methods**). Similarly, we found no difference in the gene family composition of samples between cohorts (PERMANOVA on Jaccard dissimilarity, p> 0.05; **Supplementary** Figure 4a), though we did observe the expected relationship between overall gene family composition and the phylogroup membership of the most abundant *E. coli* strain (PERMANOVA, p< 0.05; **Supplementary** Figure 4b).

### Type 1 pili are not differentially expressed in the feces of healthy women vs. those with rUTI

The lack of gene content differences between cohorts extended to well-studied urovirulence factors, including type 1 pili (T1P). T1P bind to and aid in invasion of bladder epithelial cells during UTI and support *E. coli’s* colonization of the gut ^30–32^. T1P expression is regulated, in part, by the orientation of *fimS*, an invertible, non-coding promoter element directly upstream of the *fim* operon that modulates *fim* transcription^33^. The operon encoding T1P, *fim*, was detected in all but one sample across the enriched metagenomes (**Supplementary Table 10**), as expected from previous analysis of sequenced isolates^34–36^. We also observed the non-coding *fimS* sequence in 178 of 188 (95%) enriched metagenomes despite it not being included in our probe design (**Supplementary** Figure 5-6a). From the 140 samples having at least five reads extending into the *fimS* region, which enabled statistical evaluation of whether *fimS* was in the ON (enabling *fim* expression) or OFF (inhibiting *fim* expression) orientation **(Supplementary Table 7, Methods),** we determined that only 7.7% ± 1.9% of *fimS* in each sample was in the ON orientation, regardless of rUTI history (**Supplementary** Figure 6b). However, there was considerable inter-sample variation (**Supplementary** Figure 6b), with some samples having >10% of *fim* in the ON orientation, and some having none. For samples with more than 60,000 RNA and DNA *E. coli* reads post-HS **(Supplementary Table 7, Methods**), we confirmed that *fimS* orientation was proportional to expression of the downstream *fim* operon (p=0.0004; **Methods**), as expected ^33,37,38^.

### Broad shift towards expression of aerobic metabolism genes in the rUTI gut

Given that UPEC’s transcriptional state leading up to entry into the urinary tract has been shown to be more predictive of mouse bladder colonization than its gene content ^10^, we explored expression-level differences between healthy and recurrer gut *E. coli* (**Supplementary Table 7**, **Methods**). We identified a set of 2,182 expressed genes with evidence for conservation in both cohorts, four of which were significantly differentially expressed (**Supplementary Table 11, Figure 4a, Methods**). A gene encoding a suspected exporter of dipeptides and arabinose, *ydeE*, was significantly over-expressed in samples from the recurrer cohort (p=0.048), while genes encoding L-fucose isomerase (*fucI*), aspartate ammonia lyase (*aspA*), and a subunit of the fumarate reductase complex (*frdA*), were all significantly under-expressed (all p=0.035). *dcuA* (p=0.054), encoding a C4-dicarboxylate transporter capable of exchanging fumarate for succinate, and *fucO* (p=0.095), encoding lactaldehyde reductase, were also under-expressed in recurrer gut *E. coli* . An analysis of just samples with the same *E. coli* phylogroup showed similar trends (**Supplementary Results C, Supplementary** Figure 9**, Supplementary Table 14)**.

**Figure 4.**
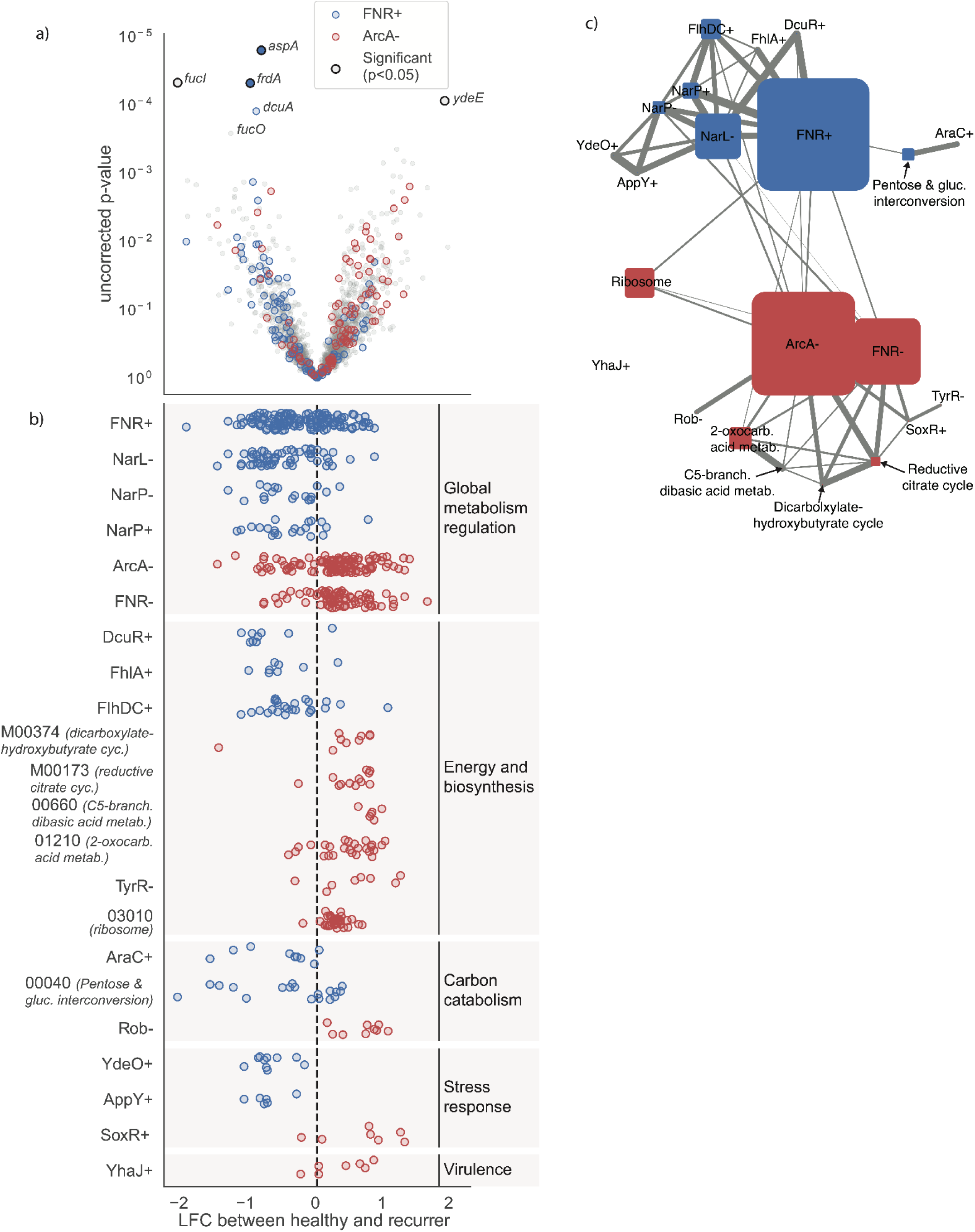
Shift towards aerobic metabolism in the rUTI gut. a) Log fold change (x-axis) and significance (y-axis) of the differential expression (DE) of 2,182 *E. coli* genes between stool from healthy and rUTI women. The 4 individual genes that were significantly DE after false discovery rate correction are indicated with black outlines and labeled. *dcuA*, which was near the significance threshold, is also labeled. Genes are colored by inclusion in the global FNR+ and ArcA- regulons. b) Distribution of fold-changes for genes in each of the 22 gene sets (KEGG pathways/modules or regulons) significantly enriched among under- or over-expressed genes (**Supplementary Table 12**). Gene sets are colored by enrichment among over- (red) or under-expressed genes (blue) and are grouped by metabolic function. c) Network diagram showing the interconnectedness of the DE gene sets, colored according to up- or down-regulation as in b).

Some of the genes found to be under-expressed in recurrer gut *E. coli* are known to work together to carry out either anaerobic respiration on fumarate (*frdA*, *aspA*, *dcuA*), or anaerobic fermentation of fucose (*fucI, fucO*) (**Supplementary** Figure 8). To conduct a formal pathway analysis, we grouped genes into transcription factor activator and repressor (noted by “+” or “-”) regulons ^39,40^, and KEGG Pathways and Modules ^41^. Roughly two-thirds (1,417 of 2,182) of the genes included in differential expression (DE) testing belonged to at least one of 284 gene sets. Using Gene Set Enrichment Analysis (GSEA) ^42^, we determined that 22 gene sets were over-represented among genes either over-expressed or under-expressed (11 each) in the recurrer cohort (p<0.05) (**Supplementary Table 12**, **Figure 4b-c**). These included the global repressor of aerobic metabolism, ArcA- (enriched among over-expressed genes), the global activator of anaerobic metabolism, FNR+ (enriched among under-expressed genes), and the global co-regulators of nitrate response, NarL- and NarP± (enriched among under-expressed genes). NarL/P and FNR co-regulate a large number of genes involved in anaerobic respiration on fumarate, including several that were also identified in the individual gene-based analysis (**Supplementary Table 11**). Of the 332 genes that were part of at least one enriched gene set, 70% (233 of 332) were regulated directly by ArcA, FNR, or NarL/P.

The other 16 enriched gene sets were smaller, largely overlapped with the global regulons (**Figure 4c**), and helped to specify differences in *E. coli* behavior between these two gut habitats. Gene sets over-expressed in the recurrer cohort pointed to increased expression of: (i) TCA cycle enzymes (KEGG M00347, dicarboxylate-hydroxybutyrate cycle; KEGG M00173, reductive citrate cycle); (ii) amino acid biosynthetic machinery (KEGG 00660, C5-branched dibasic acid metabolism; KEGG 01210, 2-oxocarboxylic acid metabolism; TyrR-); (iii) *β*-oxidation of fatty acids (Rob-); (iv) reactive oxygen and nitrogen species (RONS) detoxification (SoxR+); (v) ribosomes (KEGG 03010); and (vi) the YhaJ+ regulon, which may have strain-specific roles in virulence and is entirely unrelated to the ArcA regulatory network ^43^. Under-expressed genes in the recurrer cohort pointed to decreased expression of: (i) enzymes for anaerobic respiration on fumarate (DcuR+, FhlA+, FlhDC+); (ii) high affinity oxygen cytochromes, potentially used for redox balancing during fermentation (YdeO+, AppY+); and (iii) arabinose and xylose catabolic enzymes in pathways mostly unrelated to the FNR:NarL/P regulatory network (KEGG 00040, pentose/glucuronate interconversion; AraC+).

Collectively, the differences in *E. coli* gene expression suggested a broad transcriptional response to increased levels of oxygen (ArcA, FNR), nitrate (NarL/P), and reactive oxygen and nitrogen species, or RONS (SoxR), in the rUTI gut (**Figure 5**). RONS, produced during intestinal inflammation, have been shown to degrade to oxygen and nitrate in the gut, providing a growth advantage to facultative anaerobes, like *E. coli*, leading to proteobacterial blooms ^44,45^. Given the evidence for RONS-associated oxygen and nitrate in the rUTI gut and the increased expression of translational machinery, we hypothesized that *E. coli* may be growing at a faster rate in the rUTI gut than in the healthy gut. Using SMEG ^46^, which determines the difference in coverage between the origin and terminus of replication as a proxy for growth rate^46^, we observed no significant difference in estimated growth rate between the cohorts (p=0.54), with a greater average growth rate in the healthy cohort (**Supplementary** Figure 10).

**Figure 5.**
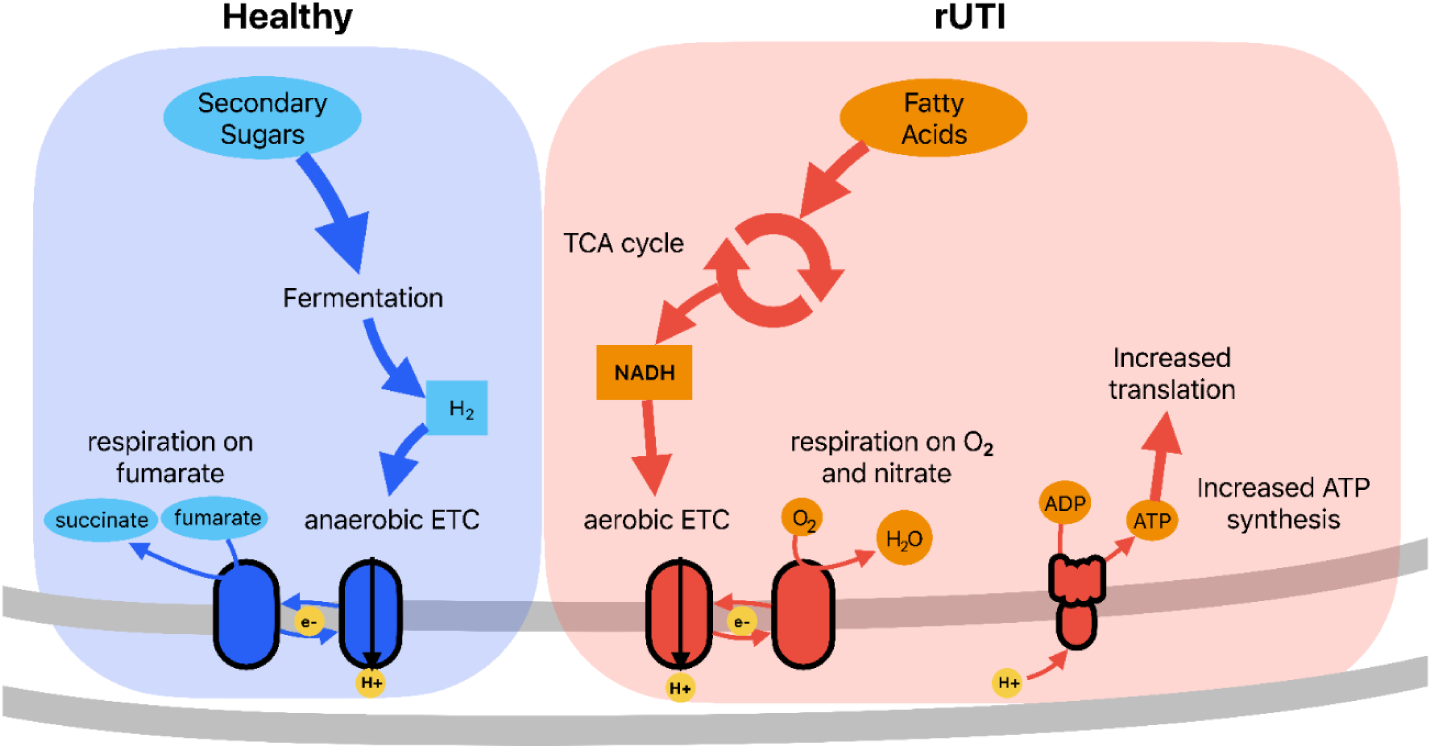
Overview of transcriptional responses that typify *E. coli* in healthy (blue) versus rUTI (red) guts.

## Discussion

Here, we provide a framework for enriching nucleic acids from a low abundance bacterial species within a complex community, where the diversity and genetic content of the species in the community are unknown *a priori* but expected to be diverse. Focusing on a low-abundance but important human commensal and opportunistic pathogen, we used this framework to develop the *E. coli* PanSelect probe set to cover the large pangenome of *E. coli*. Applied to sequencing libraries generated from a controlled mock community sample and human stool, *E. coli* PanSelect increased the coverage of *E. coli* by many orders of magnitude, equivalent to sequencing unenriched libraries to a depth required to cover the human genome 150 times. This significant enrichment in sequencing data did not skew either the *E. coli* or non-*E. coli* data fractions and revealed new fundamental details about the genetics and transcriptional programming of *E. coli* in their native gut habitat, including the dysbiotic rUTI gut.

The gut is an important pre-infection reservoir for UPEC, and it is hypothesized that conditions in the gut may modulate recurrent infection in the bladder ^47^. In an *E. coli* PanSelect-powered analysis of *E. coli* transcription in stool from women with and without a history of rUTI, we observed significant differential expression of genes encoding aspects of core energy metabolism, primarily under the control of two global regulators of anaerobic metabolism, ArcA and FNR (**Figure 4a**). A facultative anaerobe, *E. coli* carries cellular machinery for respiration on multiple electron acceptors, but will shift its metabolism towards anaerobic respiration on nitrate or aerobic respiration on oxygen whenever possible ^48,49^. In healthy conditions, the gut environment is typically deficient in nitrate and oxygen, and colonized by fermentative, obligate anaerobes^45^. However, during periods of inflammation, the intestinal epithelium produces antimicrobial reactive oxygen and nitrogen species (RONS)^45^. Facultative anaerobes like *E. coli* are more resistant to oxidative stress and can co-opt secondary products of RONS as terminal electron acceptors for respiration, resulting in changes to microbiome composition, including increased number of facultative anaerobes ^45^. In the recurrer cohort, we observed a transcriptional response consistent with increased RONS and related electron acceptors, including changes to global respiratory metabolism (ArcA, FNR) and nitrate response (NarL, NarP), and increased expression of nitric oxide and superoxide stress responses (SoxR). However, we did not observe higher relative abundances of *E. coli* in rUTI guts in our prior analysis of unenriched stool metagenomes^27^, nor increased growth rates within the HS-enriched dataset. Possibly, the growth potential provided by RONS was small enough that it was only be detected at the transcriptional level, or repeated antibiotic exposure among rUTI women may have curbed the expansion of drug susceptible strains.

A RONS-driven change in *E. coli* transcription is consistent with our prior finding of a depletion of butyrate-producing taxa in the rUTI gut ^27^. Butyrate, a short chain fatty acid (SCFA) produced by members of the gut microbiome, activates anti-inflammatory PPAR-*γ* signaling by the intestinal epithelium^50–52^. In the absence of PPAR-*γ* signaling, luminal nitrate and oxygen levels are elevated by the induction of nitric oxide synthase and the reduced expression of the β-oxidation pathway within epithelial cells, respectively ^53^.

Depletion of butyrate-producing taxa by antibiotic treatment is sufficient to disrupt PPAR-*γ* signaling and provide a growth advantage to *E. coli* ^53^. In our HS enriched dataset, apparent differences in carbon utilization between cohorts suggested further disruption to microbe-driven PPAR-*γ* signaling. Independent of the main FNR regulatory network, genes involved in catabolism of arabinose and fucose were under-expressed in the guts of recurrers. Prior work has shown that both sugars are fermented to SCFAs by the gut microbiota, with the majority of carbon in secreted fucose returned to the host as butyrate ^54–56^. Diet, the primary determinant of arabinose concentration^57^, was not different between cohorts, and fucose-liberating taxa, like *B. thetaiotaomicron*, were not depleted in the rUTI gut ^27^, suggesting that the difference in carbohydrate metabolism may be part of the broader metabolic shift induced by the gut environment.

What does the recurrer *E. coli* transcriptional program tell us about rUTI susceptibility? Transcriptional regulation is thought to be central to UPEC virulence. In a comparison of diverse urine-associated *E. coli* strains, Schreiber *et al.* found that transcriptional state, including of type 1 pili, in cultures used for bladder inoculation was a better predictor of murine urinary tract colonization than strain genetic background ^10^. Though we did not observe a difference in T1P expression between cohorts, the metabolic functions for which we observed the greatest evidence for increased activity in the rUTI gut, respiration on oxygen and nitrate, have been previously identified as key colonization factors of the bladder, including of bladder-associated intracellular bacterial communities, and the wider urinary tract ^58–61^. Up-regulation of these transcriptional programs pre-infection may make successful bladder colonization more likely. However, other inflammatory diseases of the gut, such as IBD, also lead to conditions that benefit respiring *E. coli*, but are not associated with an increased rUTI risk, to our knowledge^45,62^. Further investigations into the differences between the conditions that *E. coli* experiences in the rUTI gut and that of other inflamed gut environments may clarify the relationship between gut environment, *E. coli* lifestyle, and recurrence.

There were limitations in the design of the E. coli PanSelect probe set, and the degree to which we were able to use it to study UPEC biology within the gut microbiome. First, due to the computational complexity of producing HS probes, we adopted a coding sequence-focused design and prioritized common gene families identified through a comparative genomics workflow. Despite excluding rare gene families, we still predicted enrichment of more than 97% of the pangenome *in silico*, and observed remarkably even enrichment of *E. coli* on both a mock community and human stool samples. Enrichment extended to non-coding regions not included in probe design, such as *fimS,* due to the length of DNA fragments captured in our hybridization steps. Second, we found that enrichment of DNA-based libraries was greater than enrichment of RNA-Seq libraries, possibly due to cDNA libraries being processed in larger pools (24 vs 8 sample pools), which may have saturated probe occupancy. Finally, while not a limitation of *E. coli* PanSelect tool, our use of human stool did limit the interpretation of our data as stool is only a portion of the gut environment inhabited by *E. coli.* Stool microbial communities may not accurately reflect the gene expression profiles of *E. coli* that are adherent to the mucosa (or within the colonic crypts) or found in the small intestinal tract. Therefore, the use of stool may explain our finding that the *fimS* region was primarily in the OFF orientation, despite the importance of T1P in mediating gut colonization^30,36^. Previously, we showed that *E. coli* that are planktonic in urine have lower T1P expression (and more *fimS* in the OFF orientation) than when the same *E. coli* is adherent to the bladder mucosa^63^. Here, it is possible that *E. coli* found in stool exhibit T1P expression patterns more like planktonic urine growth, while *E. coli* that are closer to the mucosa have a different T1P expression profile. In future work, *E. coli* PanSelect, which can be applied to sequencing libraries regardless of specimen type used to construct the libraries, will be a useful tool to measure gene expression profiles of *E. coli* throughout the gastrointestinal tract, including in mucosa-associated microhabitats.

Although we focused on *E. coli* within the gut, broader applications of *E. coli* PanSelect include detection of low-abundance strains across other environments such as the skin ^64^, catheter, bladder ^7,65,66^, hospitals, ^67^ and in the food industry ^68^. Future studies may also make use of the background metagenome and metatranscriptome data, which we confirmed to be minimally biased in this work, thus, integrating the tool into established analytical pipelines and precluding the need for additional sequencing costs. In addition to applications to *E. coli,* our pangenome-based approach for probe design can be adapted into a cost-effective alternative to ultra-deep metagenomic sequencing to examine other low-abundance, genetically diverse bacteria, particularly those with numerous representative reference genomes.

## Methods

### Probe design and synthesis

#### E. coli reference genome selection, clustering, and annotation

All *E. coli* and *Shigella* (hereafter collectively referred to as *E. coli*) complete genomes were downloaded from NCBI RefSeq as of June 2017 (total of 295 genomes). To supplement the Refseq collection with additional diverse genomes, 3,141 publicly available, high quality (L50 < 20) genomes of *E. coli* that were listed in the NCBI Pathogen Detection database were downloaded from GenBank from July to August 2017 (**Supplementary Table 1**). In order to remove nearly identical Genbank genomes, we performed k-mer based clustering. All 3,141 Genbank genomes were k-merized (using 23mers) with the StrainGST “kmerize” tool from StrainGE^26^, then their pairwise all-vs-all Jaccard similarities were calculated. Single-linkage clustering was performed on these similarities at a 95% threshold to construct 1,485 genome clusters, of which 1,124 contained a single genome and 361 contained two or more Genbank genomes. Of the 361 multi-genome clusters, 67 included a genome identical to one of the RefSeq references, and were discarded. For the remaining 294, the largest reference was chosen as a representative for the cluster. Our final set included these 294 representatives, the 1,124 singletons, and the 295 RefSeq genomes. This final set of 1,713 genomes represented a large, diverse collection, with references from all eight major phylogroups of *E. coli* (**Supplementary Table 2; Supplementary** Figure 1), as determined by the tool ClermonTyping ^69^, and 515 distinct multi-locus sequence types, as determined by the tool mlst (https://github.com/tseemann/mlst).

The 1,713 genomes were then uniformly re-annotated with the Broad Institute prokaryotic genome pipeline^10,70^. Genes were clustered into orthogroups using SynerClust^71^, which resulted in a total of 174,584 orthogroups containing 8,334,026 total genes. As the computational time to analyze all these orthogroups using CATCH was prohibitive, we filtered out rare orthogroups found in fewer than three genomes, leaving 64,146 orthogroups (containing a total of 8,165,358 genes). In order to ensure that our set contained all potential instances of key genes important in clinically relevant *E. coli*, we retained all instances of orthogroups containing 59 Pfam domains of interest, obtained from a curated list (**Supplementary Table 13**). Using this list, we added back a total of 2,434 orthogroups (3,479 genes) that were found in fewer than three genomes. Our final set contained 64,580 orthogroups comprising 8,168,837 genes. In order to reduce design constraints in CATCH, thereby decreasing computational cost and time, we further clustered each orthogroup using UCLUST^72^, with an 80% identity threshold. This generated one or more clusters of genes within each orthogroup, in which all cluster members had ≥80% identity to one other. This generated 87,218 gene clusters from the 64,580 orthogroups. These gene clusters were the input for CATCH probe design.

#### Probe design and filtering

CATCH^25^ was run to generate probes for each gene cluster using the following parameters: 2 bp mismatch allowed; 25 bp cover extension; expand “N” to ACGT; 30 bp island of exact match; 60-75 bp length. In addition, three non-*E. coli Enterobacteriaceae* assemblies were used as part of the CATCH algorithm to “blacklist” probes that matched off-target sequences with a mismatch tolerance of 8 bp: a *Citrobacter ,* a *Salmonella*, and a *Klebsiella* genome (Genbank accessions GCA_000648515.1, GCF_000195995.1, and GCA_000240185.2, respectively). Representatives from these three genera were chosen as they represent well-characterized *Enterobacteriaceae*, closely related to *E. coli*; blacklisting them was thought to help to improve the specificity of the probe set to *E. coli* vs. other similar organisms. Duplicate probes were removed, resulting in a total of 911,618 unique probe sequences.

We also used an additional set of filters to remove remaining probes that might capture off-target sequences. We used the blastn tool from BLAST+^73,74^ to search probe sequences for homology against the NCBI prokaryote reference genome database (downloaded in October 2017), using the following parameters: max_target_seqs 30; evalue 1e-5; qcov_hsp_perc 80; perc_identity 80. Using these results, we removed probes that had matches of 65bp or more to: 1) ≥100 references in the database (1,798 probes removed); 2) *Bacteroidetes* references (2,470 probes removed); 3) *Firmicutes* references (14,935 probes removed). We were left with a final total of 892,415 probes that were unlikely to hit other commonly found bacterial species in the human gut.

#### In silico probe set validation

To verify that our probe set would actually be able to capture the vast majority of genes in the *E. coli* pangenome, we used blastn from BLAST+ to query our probe sequences against the entire pangenome from our set of 1,713 references, which included genes that had previously been filtered out at the probe design stage. We used probe sequences as queries for blastn with the following parameters: max_target_seqs 30; e-value 1e-5; qcov_hsp_perc 80; perc_identity 80. We retained alignments with >65 bp length and no more than 8 mismatches in the entire alignment. The probe set was considered to capture a gene if one or more probes met these criteria for a given gene.

#### Probe synthesis

For this study, the probes were synthesized by Roche, though the probe set was not specifically tailored to their technology and could be synthesized by other manufacturers. All probes could be synthesized, although 330,387 (37%) probes had one or more bases truncated from the 3’ end. The average number of bases trimmed per probe was 1.27 ± 2.16. Only 5,423 probes had 10 or more bases trimmed. All of the most highly truncated probes had low nucleotide complexity, primarily due to long stretches of homopolymers. As these changes were unlikely to affect the performance of the probe set as a whole, we used this slightly modified probe set in our experiments. The average probe length after synthesis was 73.7 ± 2.2 bp.

### Analysis of four-strain *E. coli* mock community

#### Library construction and sequencing

We used a previously reported mock community ^26^, which included 99% human DNA and 1% *E. coli* DNA. The *E. coli* DNA was composed of: i) H10407 (phylogroup A; 80%), ii) E24337A (phylogroup B1; 15%), UTI89 (phylogroup B2; 4.9%), and Sakai (phylogroup E; 0.1%). The Nextera XT library construction kit (Illumina) was used to generate sequencing libraries following the manufacturer’s recommended protocol. To enrich *E. coli* sequences in the mock library (∼100 ng into the HS reaction), we performed HS using a Roche SeqCap EZ Hypercap kit with our designed custom capture probe set. Hybridization and target capture followed the SeqCap kit instructions except that we diluted the probe pool 1:2 before use, and substituted custom Nextera adapter blocking oligonucleotides^25^ for the SeqCap HE Universal adapter and index blocking oligonucleotides. After hybridization (18 h), bead capture and washes, we performed 10 cycles of PCR with generic universal Illumina P7 and P5 primers. The final libraries were quantified by Qubit fluorometry (Thermo Fisher Scientific), and the size distribution was analyzed by TapeStation electrophoresis (Agilent) prior to Illumina sequencing. Then, pre- and post-HS libraries were sequenced on an Illumina HiSeqX, generating 21,460,598 and 75,576,717 paired-end 151 bp reads for the pre- and post-HS libraries, respectively.

#### Downsampling and quality control

Pre-HS and HS libraries were downsampled to equal depth (3 Gb, or approximately 20,000,000 paired-end reads) with Picard-Tools prior to analysis (https://broadinstitute.github.io/picard/). Quality control was performed with FastQC ^75^ and MultiQC ^76^. Due to observed heightened rates of PCR duplications in the HS libraries, both HS and pre-HS sequencing datasets were *de novo* deduplicated with FastUniq (**Supplementary Results A, Supplementary Table 3)** ^77^.

#### Calculation of enrichment, bias, and coverage levels

We assessed the enrichment of total *E. coli* within the metagenome with a one-sample t-test of the log2 fold-change for all four strains. In order to examine enrichment bias between different strains within a sample, we compared the ratios of the RAs of individual strains in HS metagenomes to the ratios in pre-HS metagenomes with paired t-tests. RAs and depth of coverage for each of the four strains were estimated using StrainGST^26^. We first built a StrainGST database consisting of just the reference genomes of the four strains of *E. coli* in the mock mixture–H10407, UTI89, Sakai, and E24377A–all downloaded from NCBI Genbank. Then, we k-merized both pre- and post-HS data and ran StrainGST (without k-mer fingerprinting) against the database to determine the RAs and depth of coverages for all four strains.

Coverage levels for each of the four strains were obtained by aligning downsampled and deduplicated data with Bowtie2 v. 2.3.4.3 ^78^ with default parameters to a concatenation of all four strains’ reference genomes. Only properly paired aligned reads with a minimum mapping quality (MQ) of 5 were retained with samtools (http://www.htslib.org/). This filtering was done to exclude reads and regions of the genomes where reads aligned equally well to different strains, with the goal of reducing bias in less abundant strains due to sequence homology to sequences deriving from the more abundant strains. Then, coverages of MQ≥5 reads were assessed using Bedtools (https://bedtools.readthedocs.io/en/latest/).

#### Assembly of E. coli mock community

Downsampled and deduplicated data were used to generate metagenomic assemblies. First, pre- and post-HS data were digitally normalized with the program khmer ^79^. Then, downsampled data were processed with Trim Galore (https://www.bioinformatics.babraham.ac.uk/projects/trim_galore/) to remove leftover adapter content. Then, reads were aligned to the hg38 reference using Bowtie2 v. 2.3.4.3 (with the “very sensitive” flag). Reads that did not align to the human genome were assembled with MetaSPAdes ^80^ with default parameters. Contigs and scaffolds <1 kb were removed and GAEMR (http://software.broadinstitute.org/software/gaemr/) was used to assess assembly metrics and determine the taxonomy of each remaining contig/scaffold.

We used BLAST+ to search for coverage of the strain reference genomes by the assembled *E. coli* contigs >1kb. Hits >1kb and with >90% identity were included in the final coverage calculation. To identify strain-specific genes in each strain in the mock community, we used an all vs. all BLAST+ approach to look for homologous genes between all reference genomes. Any gene that did not match a gene in any other strain (E-value <1e-10) was considered strain-specific.

#### Comparison of actual to expected probe coverages

To determine probe hybridization sites on each strain’s reference genome, all probe sequences were aligned using Bowtie2 v. 2.3.4.3 to each of the four reference genomes, individually. The intervals where probes aligned were designated as putative probe hybridization sites. Bedtools was used to calculate the probe coverage of the four reference genomes, as well as coverage of the reference genomes by the pre-HS and post-HS metagenomes.

### Sequencing of clinical stool samples

Total nucleic acid was extracted from stool metagenomes collected during the Urinary Microbiome (UMB) project, as previously described (Bioproject PRJNA400628) ^27^. The total (∼100 µL) nucleic acid from each sample was divided into equal aliquots for DNA and RNA sequencing. 191 samples were used for pre- and post-HS metagenomic (DNA) sequencing, of which 130 samples were used for pre- and post-HS metatranscriptomic (RNA) sequencing (**Supplementary Table 7**). The RNA aliquots were treated with DNase and Agencourt AMPure beads for a SPRI clean-up. The 130 non-enriched RNA libraries were sequenced with an Illumina NovaSeq, generating an average of 56.0 ± 56.7 million paired-end 151 bp reads. The 191 non-enriched DNA libraries were sequenced with a combination of Illumina HiSeq 2500 and HiSeq X10 technologies, as previously reported ^27^.

#### Enrichment and sequencing of E. coli PanSelect libraries

We enriched *E. coli* sequences from both DNA and RNA samples with multiplex solution HS using a Roche SeqCap EZ Hypercap kit with our designed custom capture probe set. DNA-Seq libraries were processed in pools of 8 libraries (∼200 ng each). RNA-Seq libraries were prepared as pools of multiplex RNAtag-Seq libraries from 24 RNA samples ^81,82^ and amplified by 14 cycles of PCR to generate at least 100 (mean 140) ng of each library pool for HS (one 24-plex pool per reaction). Hybridization and target capture were performed as for the mock community above. All post-HS libraries were run on an Illumina NovaSeq, generating an average of 9.6 ± 3.9 million paired-end 151 bp reads for post-HS DNA libraries, and 10.2 ± 10.7 million paired-end 151 bp reads for post-HS RNA libraries. Three samples for which HS DNA sequencing failed (low post-QC read depth) were removed from analysis (UMB13_09, UMB08_04, UMB24_08).

#### Quality assessment

Quality of sequencing files was assessed with FastQC and MultiQC. We did not *de novo* deduplicate DNA reads, as we observed far lower PCR duplication in the HS DNA libraries than for the mock community (**Supplementary Results A, Supplementary Table 6**). We processed DNA and RNA data with KneadData (https://huttenhower.sph.harvard.edu/kneaddata/) to remove low quality sequence, adapter content, and human contamination.

### Benchmarking *E. coli* PanSelect enrichment using stool samples

#### Enrichment estimation

For both DNA and RNA, per-sample *E. coli* enrichment was estimated as the fold change in the depth-normalized read coverage of the UTI89 reference genome between pre-HS and post-HS sample pairs. Global alignments (with Bowtie2 v. 2.3.4.3) were used for estimating fold change in reference coverage. Strain composition was estimated with StrainGST. To track changes to strain relative abundances with HS, strain calls were paired between pre- and post-HS samples. First, pre- and post-HS strains assigned to the same reference were paired (184 strain pairs). Next, strains assigned to different references of the same phylogroup were paired (14 strain pairs). There were 16 instances where one strain in one sample appeared to match two strains in the corresponding sample. Because it was unclear if this was due to a strain calling error, these 16 instances were removed from analysis. Remaining strains with no clear pair were assumed to only be detected in one sample (55 strain discoveries).

The expression levels of individual transcripts were used to illustrate RNA enrichment. First, ordinary least squares regression was used to find the average relationship between log10-transformed pre-HS and post-HS *E. coli* relative abundance. Three samples were identified as enrichment outliers via t-tests on studentized residuals and were removed (Benjamini-Hochberg FDR correction, p<0.1). Of the remaining samples, the 94 with >1 million post-QC pre- and post-HS reads were downsampled to 1 million reads. Reads were aligned to the UTI89 reference with bowtie2 and counted with FADU v.1.8^83^, run without the expectation maximization algorithm. Transcript expression levels were estimated in counts per million (CPM). Transcripts expressed below 10 CPM were classified as not detected (n.d.). For visualization, transcripts from all 94 pre-HS/post-HS sample pairs were plotted on the same axes (**Figure 3b**). To show density, clusters of transcripts expressed at similar rates were formed with hierarchical clustering.

#### Background metagenome and metatranscriptome analyses

Pre- and post-HS metagenomes and metatranscriptomes were downsampled to a maximum depth of 3.5 Gb. Taxonomic and transcriptomic profiles for all metagenomes and metatranscriptomes (pre- and post-HS) were calculated with MetaPhlAn3 and HUMAnN3 ^28^ for samples with more than 1,000,000 post-QC reads. *E. coli* and *Shigella* were removed from the MetaPhlAn RA table and the HUMAnN gene family output. Remaining values were sum normalized. Bray-Curtis dissimilarity values were calculated with the Python package *scipy* v.1.7.1 ( https://scipy.org/). PERMANOVA was implemented by the *adonis2* function from the R package *vegan* (https://cran.r-project.org/package=vegan).

#### Sequencing and assembly of E. coli isolate for benchmarking of metagenomic assembly

We sequenced a single *E. coli* isolate from Participant 8, timepoint 2, using a combination of Illumina and Oxford Nanopore Sequencing. Sequencing and hybrid assembly was performed as previously published ^84^. This assembly has been submitted to NCBI Genbank with accession GCF_011751425.1.

#### Metagenomic assembly

Metagenomic assemblies were generated for ten samples using all available sequencing data. Digital normalization, adapter trimming, assembly, and calculation of assembly metrics were performed as for the mock community data. Metrics were calculated based on only contigs assigned to *E. coli* using blastn. After assemblies were produced, a binning program, MetaBat2^85^, was used to produce metagenome-assembled genomes (MAGs). MAGs were analyzed with CheckM ^86^ to determine taxonomy and assembly completeness for MAGs that were classified as *Enterobacteriaceae* by CheckM.

### Analyses of full set of enriched stool samples

#### Comparison of gene content between cohorts

We calculated gene family presence/absence profiles for enriched metagenomes with PanPhlAn3 (v. 3.1), run with the UniRef90 *Escherichia coli* pangenome generated on Nov 2, 2020 by the Segata Lab ^28^. Samples were filtered based on evenness of *E. coli* coverage with the profiling workflow contained within PanPhlAn (*panphlan_profiling.py)*, run with the ‘very sensitive’ parameters (--min_coverage 1 --left_max 1.70 --right_min 0.30).

We used omnibus and per-gene tests to test for differences in gene content between the recurrer and healthy cohorts. First, we quantified differences in overall gene content profiles between samples with the Jaccard Index and tested for differences between cohorts with PERMANOVA. We used the *adonis2* implementation of PERMANOVA (R package *vegan*) to assess the marginal effect of cohort, controlling for subject. In order to test for differential abundance of individual gene families, we used logistic regression, and Fisher’s exact tests for genes where we had issues with model fitting. For logistic regression, we selected the genes that were found in between 10% and 90% of all samples, and detected in at least one sample from both the recurrer and healthy cohorts (4,114 gene families). We fit models with the function *pglmm* from R package *phyr* v.1.1.2^87^, using random intercepts for subjects and a phylogenetic covariance structure based on the most abundant strain identified within each sample by StrainGST. No gene families were significantly differentially abundant after Benjamini-Hochberg false discovery rate correction. Samples from individuals in the rUTI cohort who did not have a UTI during the study period (e.g. non-recurrers) were included in modeling, but only the contrast between the recurrer and healthy cohorts was reported. To evaluate the remainder of gene families for which we had issues with model fitting, we used Fisher’s exact tests to calculate the significance of the association between gene families and rUTI history, at both the sample and subject carriage levels. For the subject level, we called a gene present in a subject if it was identified in at least one sample in a subject series. For both sets of models (subject and sample level), the most significantly differentially abundant genes were those that were already shown to be insignificant by the more comprehensive logistic regression model. Thus, we concluded that the remaining genes were not differentially abundant either.

#### fimS structural variation profiling

We estimated the fraction of *fimS* in the ON orientation within enriched metagenomes by aligning reads (Bowtie2 v. 2.3.4.3; MQ>5) to a reference containing a copy of the UTI89 genome with *fimS* in the ON orientation and a copy with *fimS* in the OFF orientation. For many samples, we observed a mixture of alignments to both the ON and OFF references, indicating subpopulations of *fim-*expressing (piliated) and non-expressing (smooth) *E. coli* within the same samples. We used the proportion of reads aligning uniquely to the ON orientation as an estimate for the size of the piliated population. Because there was significant inter-sample variation in *E. coli* RA (and thus *fimS* coverage), we were able to estimate the RA of the piliated population more accurately in some samples than others. In order to reflect this variation in our estimates of average *fimS* activation, we used weighted averages and regressions with weights proportional to per-sample *fimS* coverage (ON + OFF), and filtered samples with <5 *fimS*-aligning reads from analysis (**Supplementary Table 7**).

#### Selection of E. coli coverage threshold for fim and differential expression analysis

Post-HS RNA and DNA samples were aligned to the UTI89 reference genome with bwa-mem v.0.7.17-r1188 ^88^, and alignments were counted with FADU^83^ (run with parameters: *-M -p*). We then filtered samples based upon total post-HS DNA and RNA *E. coli* content, using a threshold of 60,000 UTI89 coding sequence-aligning reads. We determined this threshold by examining the relationship between RNA *E. coli* content and observed RNA transcript diversity. We used ‘relative abundance-weighted transcript diversity’ as a metric, defined as the sum of average RAs of all unique transcripts detected in a sample. We selected our threshold of 60,000 *E. coli* reads, because unique transcripts representative of 80% of the average *E. coli* transcriptome could be observed in samples with >60,000 *E. coli* reads (**Supplementary** Figure 7).

#### Relationship between fimS orientation and fim operon expression

For testing the relationship between *fimS* orientation and *fim* operon expression, we used the subset of samples for which we could both estimate *fimS* orientation (5 *fimS*-aligning reads) and measure gene coverage and expression (60,000 *E. coli* DNA and RNA reads) (**Supplementary Table 7**) We quantified *fim* operon coverage and expression as the sum of coverage of all genes in the protein coding *fim* operon (*fimAICDFGH*). We used mixed effects models for metagenome-controlled differential expression testing, as described in *Zhang et al 2021* ^89^. Models were fit with the form: *fim RNA(log cpm) ∼ fim DNA(log cpm) + (1|sunject)* with the function *lme* from the R package *nlme*. To control for variation in *E. coli* content between samples, we used weights proportional to sample *E. coli* content: *weights=varFixed(∼ (1/DNA_ecoli + 1/RNA_ecoli))*.

#### Selection of samples and genes for differential expression testing

For differential expression testing between cohorts, we filtered samples based upon total coverage of *E. coli* (60,000 DNA and RNA reads, see above) and coverage evenness, as determined by the profiling workflow contained within PanPhlAn3, run with ‘very sensitive’ parameters (--min_coverage 1 --left_max 1.70 --right_min 0.30) on alignments to the UTI89 reference (bwa-mem v.0.7.17-r1188, counted with FADU, run with parameters *-M -p*; see above) (**Supplementary Table 7)**. We used genes that were classified as ‘single copy’ within metagenomes by PanPhlAn. For differential expression testing, we selected a subset of genes present in at least 50% of all metagenomes from both the recurrer and healthy cohorts (15 samples/cohort) and expressed in at least 15 metagenome/metatranscriptome pairs across the full study. Gene presence was defined as coverage by at least 20 reads within a metagenome. Expression was defined as presence with additional coverage by at least 20 reads within the associated metatranscriptome. We set the read threshold (20) based on the *E. coli* content range of samples included in DE testing. The sample with the 15th greatest RNA *E. coli* content had 10-fold more *E. coli* than the sample with the least *E. coli*. Thus, setting the threshold at 20 reads ensured that for any gene classified as expressed in 15 samples, we would have sensitivity to detect at least two reads in all samples where the gene was expressed at an equal or greater rate.

#### Between cohort differential expression testing

We used mixed effects models for per-gene metagenome-controlled differential expression tests^89^. We fit models of the form: *RNA(log cpm) ∼ DNA(log cpm) + cohort + E. coli RA(log %) + (1|sunject)* in with the function *lme* from the R package *nlme*. To control for variation in *E. coli* content between samples, we used weights proportional to sample *E. coli* content: *weights=varFixed(∼ (1/DNA_ecoli + 1/RNA_ecoli)).* We included *E. coli* RA as a fixed effect based on model comparisons using Akaike’s information criterion (AIC). We quantified the coverage of genes within metagenomes and metatranscriptomes as copies per million scaled by *E. coli* content (CPM), and used a natural log transformation for variance stabilization of RNA and DNA CPM values, as well as *E. coli* RAs. RNA zeroes were replaced before transformation with a gene-specific pseudocount equal to half the lowest non-zero RPM value measured for each gene, as done in Zhang et al. (2021). There were no zero values for DNA (because we used single copy genes) or *E. coli* RA.

#### Gene set enrichment analysis

We used Gene Set Enrichment Analysis (GSEA) ^42^ to test for overrepresentation of gene sets among genes over and under-expressed in the recurrer cohort. We grouped genes into TF regulons, using gene:TF interactions reported in RegulonDB^39^, as well as KEGG Modules and Pathways. For TFs with dual activity, we grouped genes into separate regulons consisting of genes activated and repressed by the TF. Because RegulonDB reports regulatory information for *E. coli K12* and we used a *E. coli UTI89* reference, the TF:gene interactions were not immediately transferable to our dataset. We used SynerClust ^71^ to pair orthologs between the two reference genomes, and transferred annotations from the *K12* reference to UTI89 orthologs. For annotation of KEGG Pathway and Module membership, we used KEGG gene name annotations from RegulonDB and the KEGG *E. coli K12* (*eco*) Pathway and Module maps. For GSEA, we used gene sets that were five genes in size or larger. We used the t-scores from the differential expression tests (reported above) as input for GSEA. We reported GSEA results at an FDR-corrected significance threshold of 0.05.

#### Metagenomic growth rate estimation

The growth rate of *E. coli* strains within metagenomes was estimated as the difference in coverage between the origin and terminus of replication, as implemented in SMEG^46^. We constructed a SMEG species database using the strain reference genomes reported by StrainGST, and ran SMEG using the ‘reference based’ mode with the strain RA estimated by StrainGST, using the post-HS sequencing data. Per sample, we calculated the average *E. coli* growth rate as the average of strain growth rates, weighted by strain RA. We used linear mixed effects models to compare strain growth rate between cohorts, of the form *growth rate ∼ cohort + (1|sunject)*.

### Statistical analysis and graphical plotting

All statistical analysis and plotting was performed in R v3.6^90^ ggplot2^91^, data.table (https://rdatatable.gitlab.io/data.table/), Rmisc (https://cran.r-project.org/package=Rmisc). The following Python libraries were used: pandas v.1.3.2^92^, numpy v.1.21.2^93^, scipy v.1.7.1^94^, statsmodels v.0.12.1^95^, biopython v.1.7.9^96^, pysam v.0.19.1^97^, matplotlib v.3.4.3^98^, and seaborn v.0.11.2^99^.

## Data availability

Post-HS sequencing data has been submitted to SRA to NCBI’s Sequence Read Archive under Bioprojects PRJNA685748 (mock community) and PRJNA400628 (UMB stool samples). Pre-HS sequencing data was previously submitted under these same Bioprojects (pre-HS DNA data for the mock community and stool samples from the UMB project). The assembly of the *E. coli* isolate UMB08_02 has been submitted under accession GCF_011751425.1.

## Supporting information

Supplementary Results and Figures

## Acknowledgements

We would like to thank Curtis Huttenhower and members of his lab as well as members of the Bacterial Genomics group for helpful discussions. We would also like to thank Broad’s Genomic Platform and Jonathan Livny and the Microbial ‘Omics Core for their assistance with data generation.

This project has been funded with Broad NextGen funds to AME and Federal funds from the National Institute of Allergy and Infectious Diseases, National Institutes of Health, Department of Health and Human Services, under Grant Number U19AI110818 to the Broad Institute, R01AI165915 to Washington University and R01DK121822 to Washington University and the Broad Institute.

