## Supplementary Results and Figures for "Distinct *Escherichia coli* transcriptional profiles in the guts of recurrent UTI sufferers revealed by pangenome hybrid selection"

**Corresponding author

Affiliations: [1]Infectious Disease & Microbiome Program, Broad Institute, Cambridge, MA 02142, USA; [2]Delft Bioinformatics Lab, Delft University of Technology, Van Mourik Broekmanweg 6, Delft, 2628 XE, The Netherlands; [3]Department of Molecular Microbiology, Washington University School of Medicine, St. Louis, MO, USA.; [4]Center for Women's Infectious Disease Research, Washington University School of Medicine, St. Louis, MO, USA; [5]Division of Allergy and Immunology, Department of Medicine, Washington University School of Medicine, St. Louis, MO, USA

### Supplementary Results

**Index**

A. Sequence quality
B. *E. coli* PanSelect greatly increased the capability to assemble low-abundance *E. coli* genomes.
C. Differential expression testing on phylogroup-controlled subset of samples mirrored results from full analysis
D. Supplemental References
E. Supplementary Figures 1-10

#### A. Sequence quality

For both the mock community and the human stool samples, we observed similar sequence and base quality metrics in pre- and post HS libraries. GC content was slightly elevated post-HS, due to increased relative abundance of *E. coli* (**Supplementary Tables 3,6).** For human stool samples, base quality was slightly better for post-HS RNA than pre-HS RNA, but otherwise, metrics were very similar. We expected higher duplication rates in the enriched libraries because the HS library construction protocol included an additional PCR amplification step as well as amplification from specific probe sites. For the mock community samples, duplication rate increased in the HS-enriched library from 5.6% pre-enrichment to 30.6% post-enrichment (**Supplementary Table 3)**. Thus, we deduplicated mock community libraries with FastUniq [1]. For human stool data, duplication rates decreased with enrichment from 32% pre-enrichment to 24% post-enrichment, so we did not deduplicate (**Supplementary Table 6)**.

**B. *E. coli* PanSelect greatly increased the capability to assemble low-abundance *E. coli* genomes.**

Mock community. We produced metagenomic assemblies for the four strains using MetaSPAdes [2]. As expected, we were able to produce substantially more complete assemblies using the HS-enriched data (**Supplementary Table 4**). The pre-HS assembly was only 4.3 Mb in total length (less than the length of a single *E. coli* genome) and very fragmented, while the post-HS assembly was 7.85 Mb and more contiguous (including one contig nearly 60 kb in length).

In order to assess assembly of each of the four strains individually, we aligned contigs against their respective reference genomes. Before HS, 59-69% of each reference was covered; after HS, this increased to over 95% (**Supplementary Table 5**). Strikingly, the proportion of each genome assembled on long contigs (>10kb) increased from <1% pre-HS to 35-42% post-HS. The proportion of each genome assembled on long contigs was similar for all four strains (ranging from 0.01-0.8% RA) due to shared genomic content.

To assess our ability to specifically assemble genomes at different abundances, we identified regions of the genome unique to each of the four references (Methods) and examined their assembly (**Supplementary Table 5**). With *E. coli* PanSelect, the percentage of strain-specific genes assembled increased from 19% to 77% in E24377A, the lowest-abundance strain, and from 51% to 97% in H10407, the highest-abundance strain. When looking at strain-specific genes captured on contigs >10kb, the percentages increased from 0.4% to 10% for E24377A, and from 2.6% to 55.3% for H10407.

Human stool. Metagenome-assembled genomes (MAGs) produced out of enriched metagenomes were also more accurate and complete. We assembled MAGs out of ten UMB samples with roughly 1% *E. coli*, using equivalently downsampled pre-HS and HS metagenomes. All ten MAGs assembled out of enriched data were roughly the expected size of an *E. coli* genome, highly contiguous, and complete, as assessed with CheckM marker genes (**Supplementary table 8**). For an eleventh UMB sample, we compared MAGs assembled out of pre-HS and HS metagenomes to a near-complete reference assembly produced by isolating and sequencing a strain with a combination of Illumina and Oxford technologies (Methods). The MAG assembled from the HS metagenome was nearly 5 MB and covered over 99% of the isolate reference, a stark improvement over the 0.016 MB MAG assembled out of the pre-HS data **(Supplementary table 9)**.

### C. Differential expression testing on phylogroup-controlled subset of samples mirrored results from full analysis

We found no differences in phylogroup distribution between samples from women with and without rUTI history in our previously reported analysis of a larger set of unenriched UMB stool metagenomes [3]. However, among the set selected for DE testing, there was a significant difference in composition between the cohorts, with over-representation of single-strain B2 and D samples among samples from the recurrer cohort ($\chi^2$ test, p=0.034). 43% of samples contained multiple strains, with single-strain samples primarily containing phylogroups B2 (25%) and D (16%) (Supplementary Table 14 )

To confirm that our differential expression results were not driven by inter-cohort variation in phylogroup composition, we re-ran differential expression tests on the subset of samples containing a single B2 strain. On this smaller set, we were able to fit models for 659 of the 2,182 genes that were included in the full differential expression analysis. For inclusion in the B2-only analysis, we required genes to be detected in all but one metagenome from the recurrer and healthy cohorts, and expressed in at least five metatranscriptomes (using 20 reads per sample as a cutoff for detection, as done for the full analysis). Given the reduced sample size, the models fit to the B2 subset lacked statistical power and no genes were significantly differentially expressed after false discovery rate correction. However, there were significant correlations between the log fold changes estimated on the full set of samples and the B2 subset (spearman correlation, r=0.51, p=2.2e-45), and the model t-scores (r=0.52,p=2.3e-47). The concordance was especially strong for genes that were significantly differentially expressed between cohorts (**Supplementary Figure 8**). Accounting for variation in significance of differential expression, the weighted correlations between estimated log fold changes and t-scores were 0.91 and 0.90, respectively.

### E. Supplementary Figures

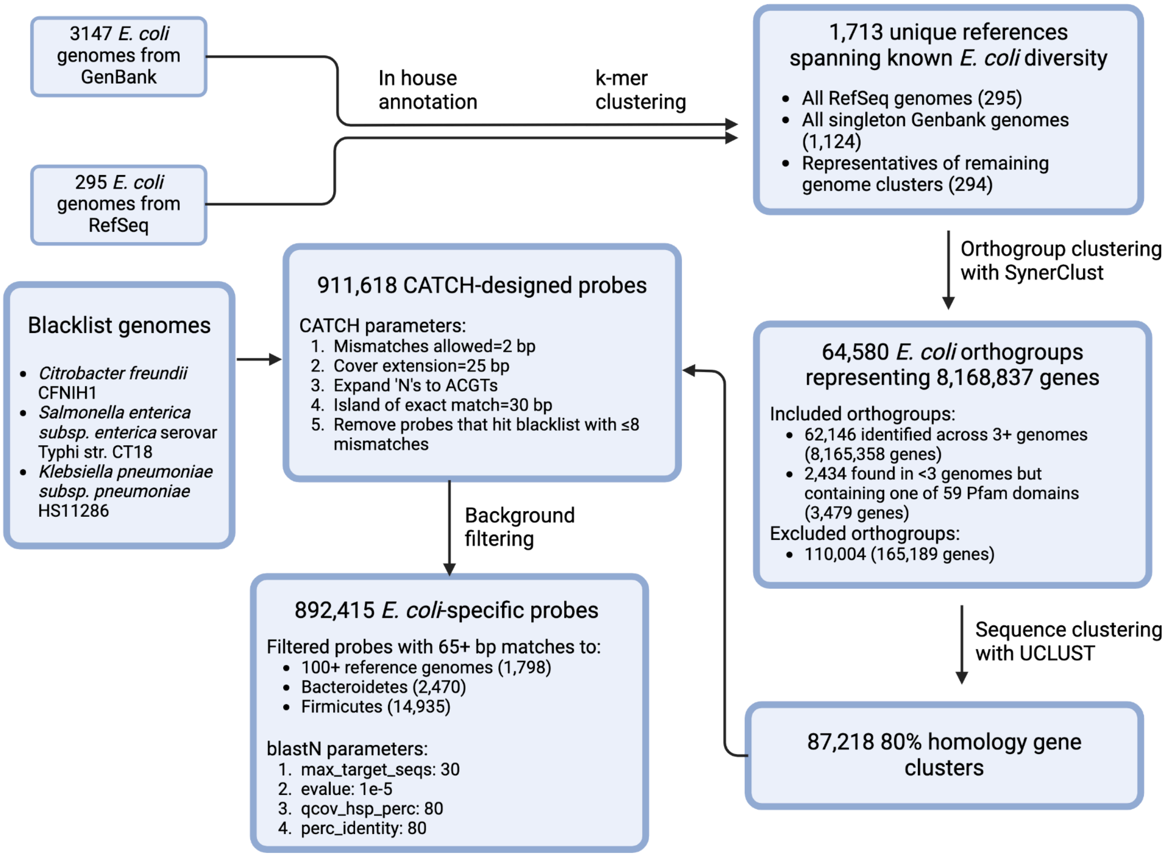


**Supplementary Figure 1. Details of probe design procedure.** A flow chart describing probe design, including parameters used, as well as the number of genomes, orthogroups, genes, and probes that were involved at each step.

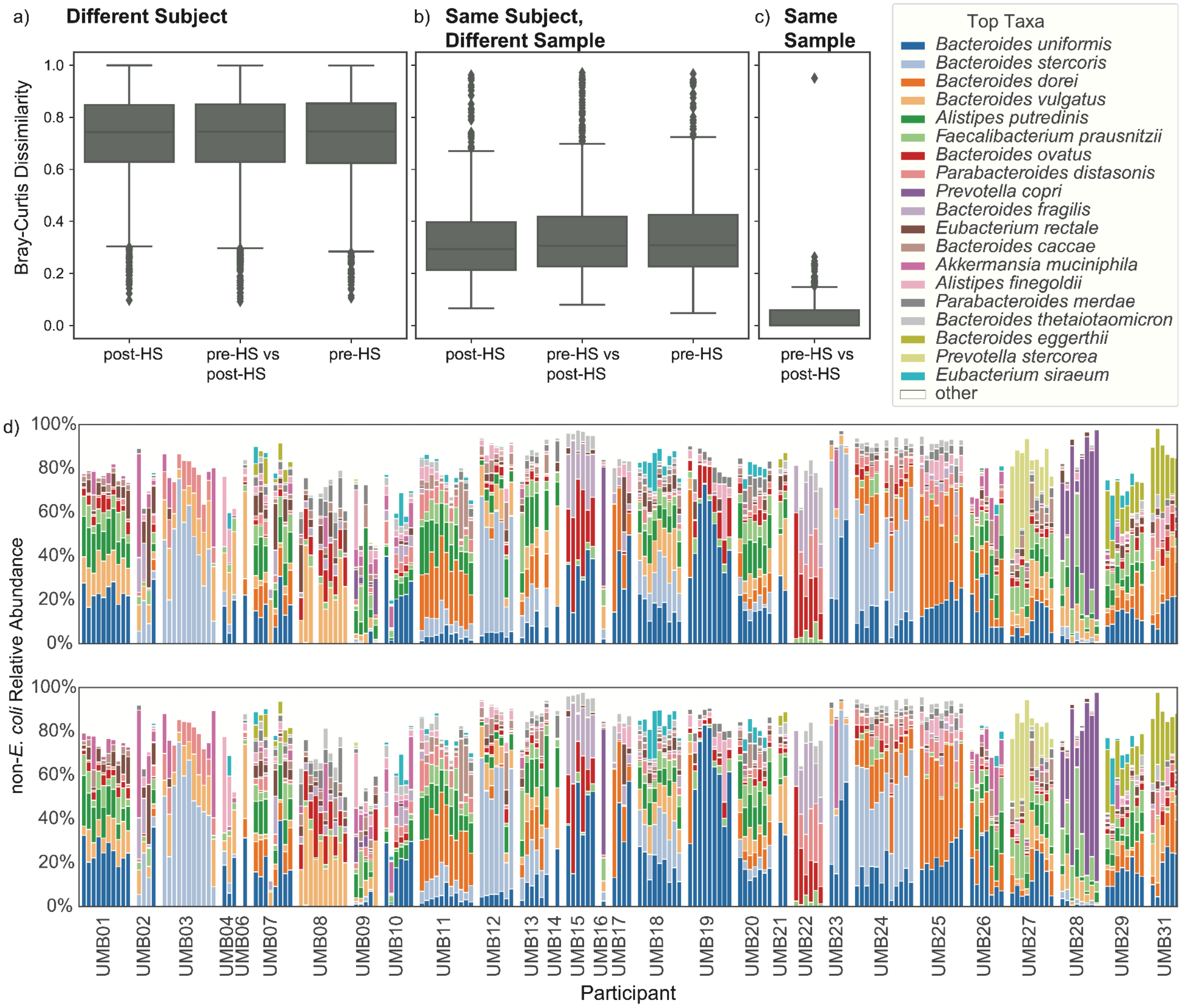


**Supplementary Figure 2. The taxonomic composition of the background metagenome is unaltered by HS.** a-c) Distributions of Bray-Curtis dissimilarities between the non-*Escherichia* (and non-*Shigella*) metagenomic fractions, separated by sequencing methodology (pre- vs pre-HS, post- vs post-HS, and pre- vs post-HS). Metagenomes were sequenced from a) samples from separate study participants, b) samples from the same participants, but at separate time points, and c) the same samples. d) Relative abundances of the most abundant taxa are plotted for each sample. Samples are sorted by subject, with post-HS on the top row and pre-HS on the bottom.

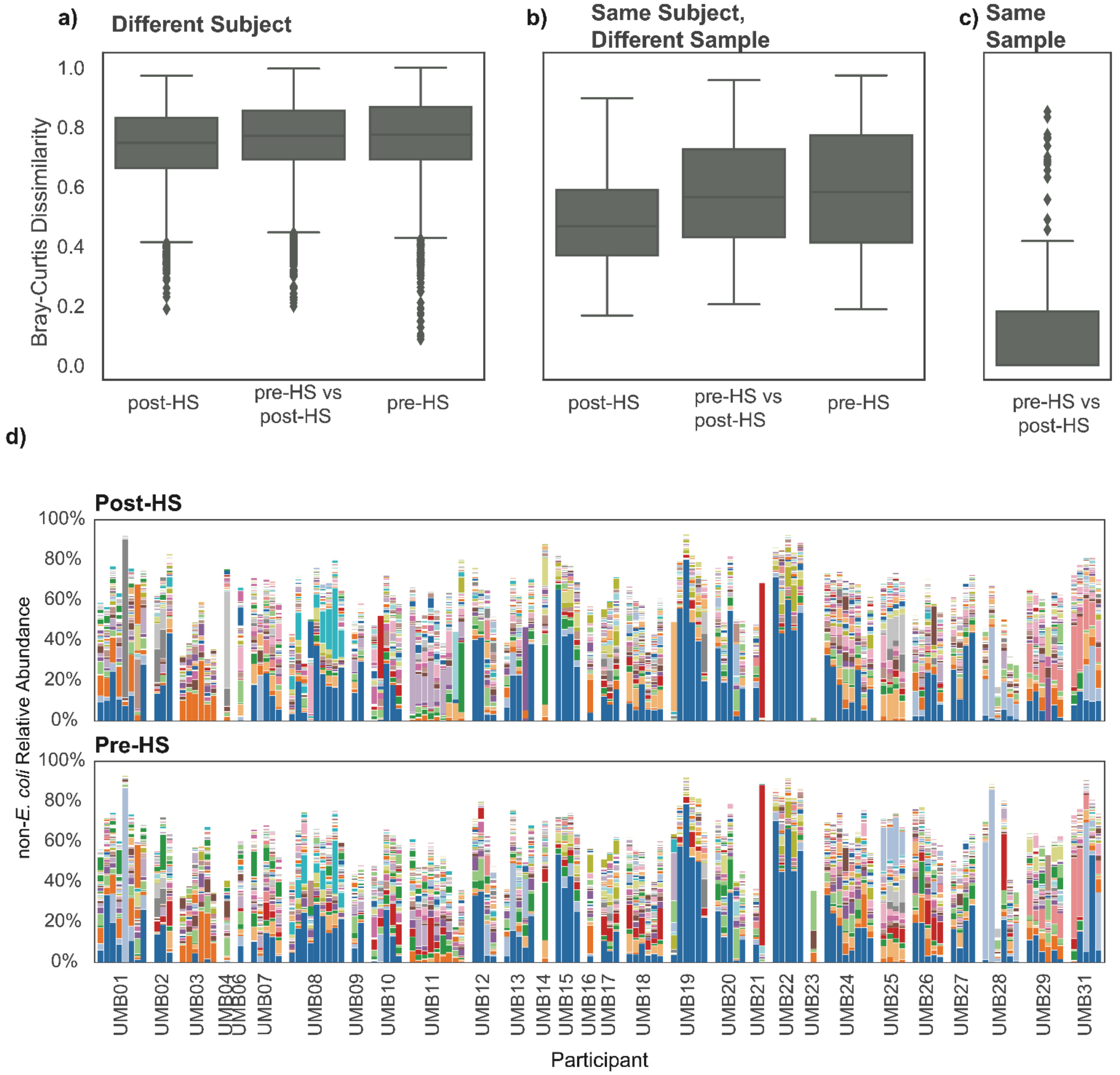


**Supplementary Figure 3. The gene family composition of the background metatranscriptome is unaltered by HS.** Gene family transcription profiles (generated with HUMAnN3) were filtered to remove transcription of *Escherichia* genes. a-c) Distributions of Bray-Curtis dissimilarities between the non-*Escherichia* transcription profiles, separated by sequencing methodology (pre- vs pre-HS, post- vs post-HS, and pre- vs post-HS). Metatranscriptomes were sequenced from a) samples from separate study participants, b) samples from the same participants, but at separate time points, and c) the same samples. d) Expression levels of the top gene families. Samples are sorted by subject, with post-HS on the top row and pre-HS on the bottom.

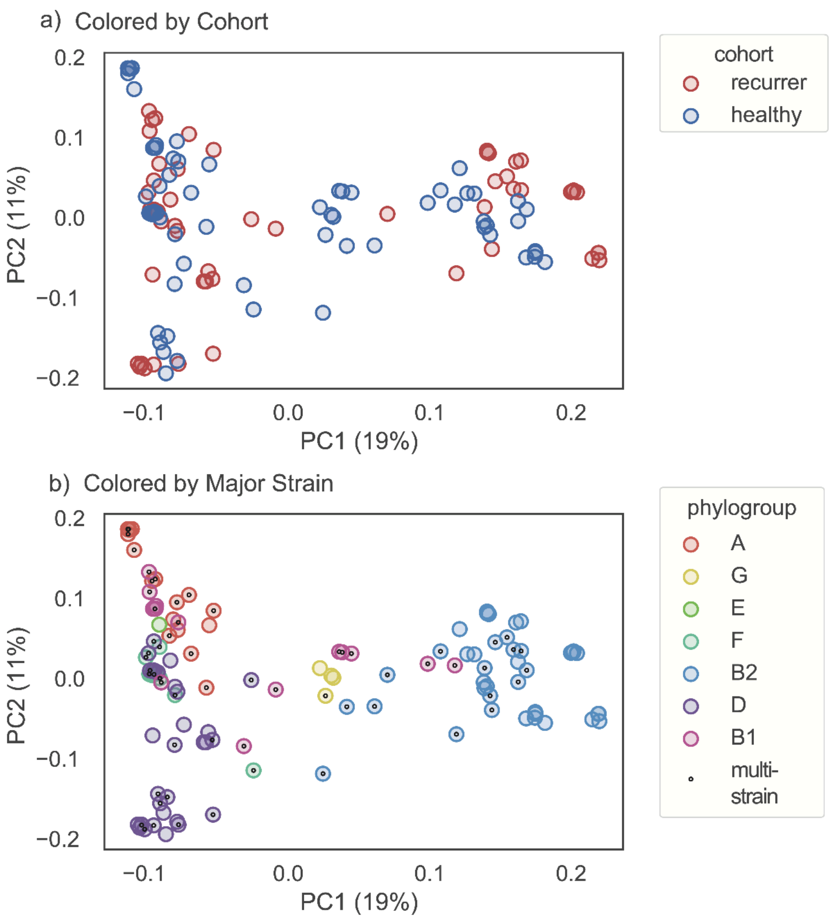


**Supplementary Figure 4. rUTI-associated *E. coli* are not characterized by unique gene content**. Principal component plots of the jaccard dissimilarities between sample pangenomes, colored by (a) cohort and (b) the phylogroup of the major strain. In b), black dots indicate multi-strain samples.

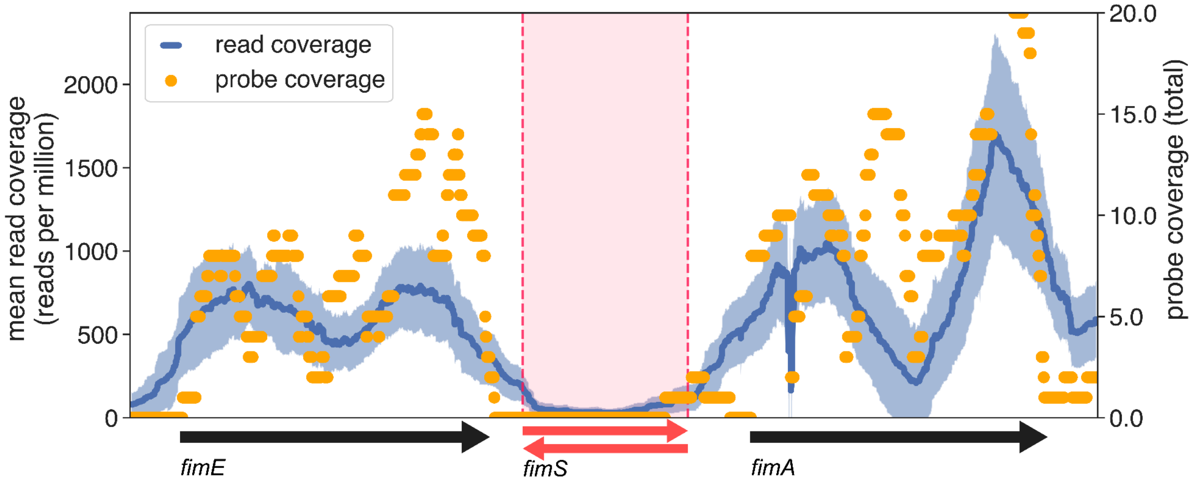


**Supplementary Figure 5. Probe coverage of neighboring genes is sufficient to enrich boundary regions of *fimS*, allowing determination of *fimS* orientation.** Mean coverage depth for the *fimS* region of all 188 HS metagenomes (blue), overlaid with total coverage by HS probes (orange). Dark blue shading indicates one standard deviation above and below the mean HS read coverage. Coverage of the genes neighboring *fimS*, *fimE* (recombinase) and *fimA* (pili structure), are also shown to scale. The Pearson correlation between mean DNA depth of coverage and per-base probe coverage is 0.77. Reads bridging from *fimE* or *fimA* into the *fimS* region (red) were used to determine *fimS* orientation.

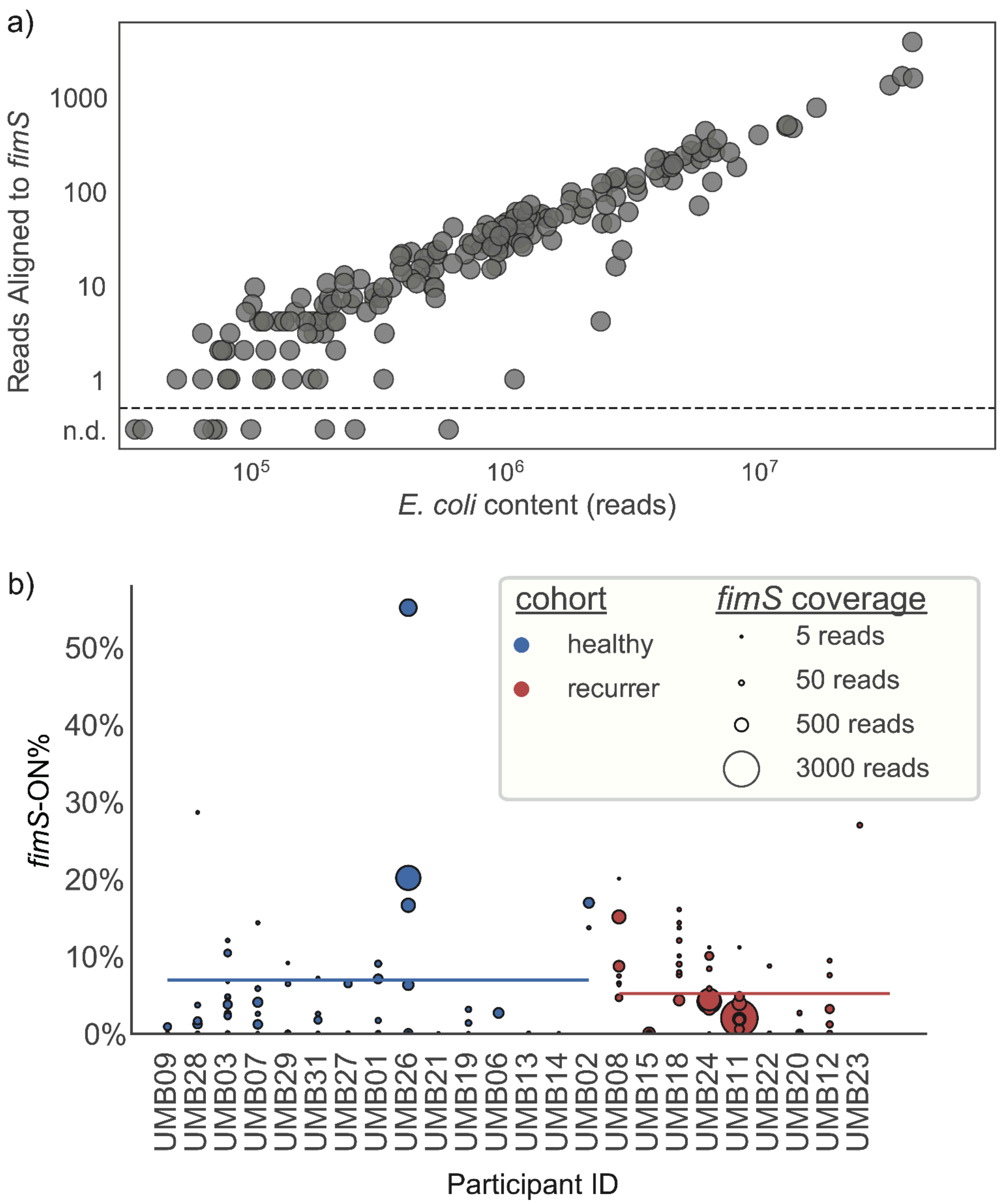


**Supplementary Figure 6. Type 1 pili regulation does not differ between healthy versus rUTI participants.** a) The number of reads aligning to *fimS* is shown, as a function of the number of total *E. coli* reads (r=0.89). Each dot represents a sample. Samples below the line marked 'n.d.' had no *fimS* detected. b) Per sample percentage of reads with *fimS* in the ON orientation, separated by cohort (blue=healthy; red=recurrer).

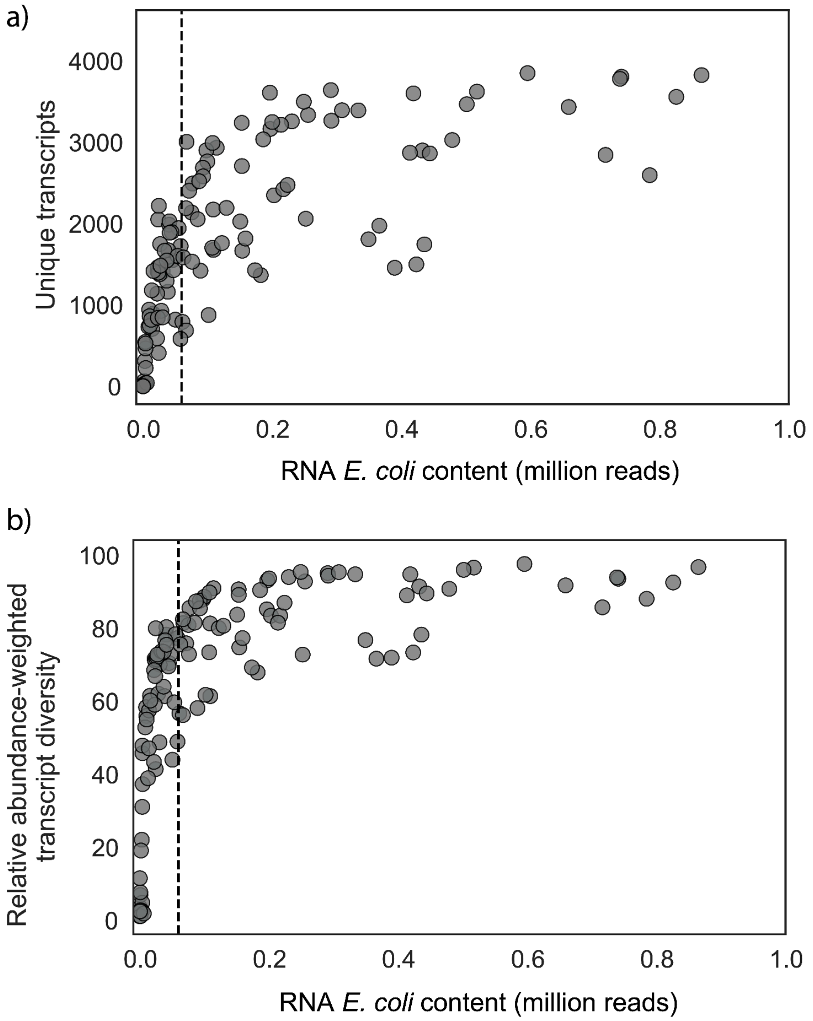


**Supplementary Figure 7. HS provided a near-saturated view of transcript diversity in samples with more than 60,000 *E. coli*-aligning RNA reads.** Samples are represented by points, with metatranscriptome *E. coli* content on the x-axis. a) An unweighted metric, the total number of unique coding sequences aligned by an RNA read, is plotted on the y-axis. b) The 'RA weighted transcript diversity', or the number of unique coding sequences observed per metatranscriptome, weighted by their average RA across the dataset, is plotted on the y-axis. Both plots are truncated to a maximum metatranscriptome *E. coli* content of 1,000,000 reads to increase resolution of the curves. Dashed vertical bars are included at the chosen threshold of 60,000 reads.

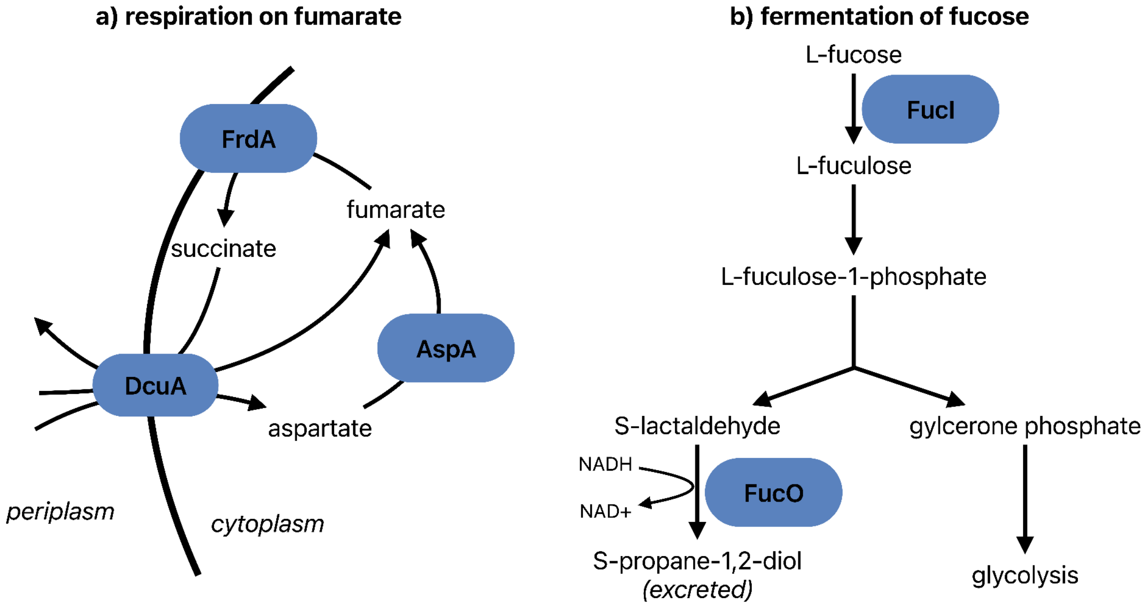


**Supplementary Figure 8. Multiple under-expressed *E. coli* genes contributed to the same pathways.** Protein names for under-expressed genes are given in blue ovals, and reactions are indicated with black arrows between compound names  a) Respiration on fumarate. The fumarate reductase flavoprotein subunit (FrdA) catalyzes electron transfer to fumarate, coupled with generation of a proton-motive force (for ATP generation). The C4-dicarboxylate transporter (DcuA) facilitates exchange of C4-dicarboxylates across the membrane, enabling uptake of fumarate and aspartate with efflux of succinate. The aspartate ammonia-lyase (AspA) catalyzes conversion of L-aspartate to fumarate and ammonia.  b) Fermentation of fucose. L-fucose isomerase (FucI) catalyzes isomerization of L-fucose to L-fuculose. Lactaldehyde reductase (FucO) catalyzes conversion of S-lactaldehyde, a byproduct of fucose degradation, to S-propane-1,2-diol, coupled with regeneration of NAD+ from NADH.

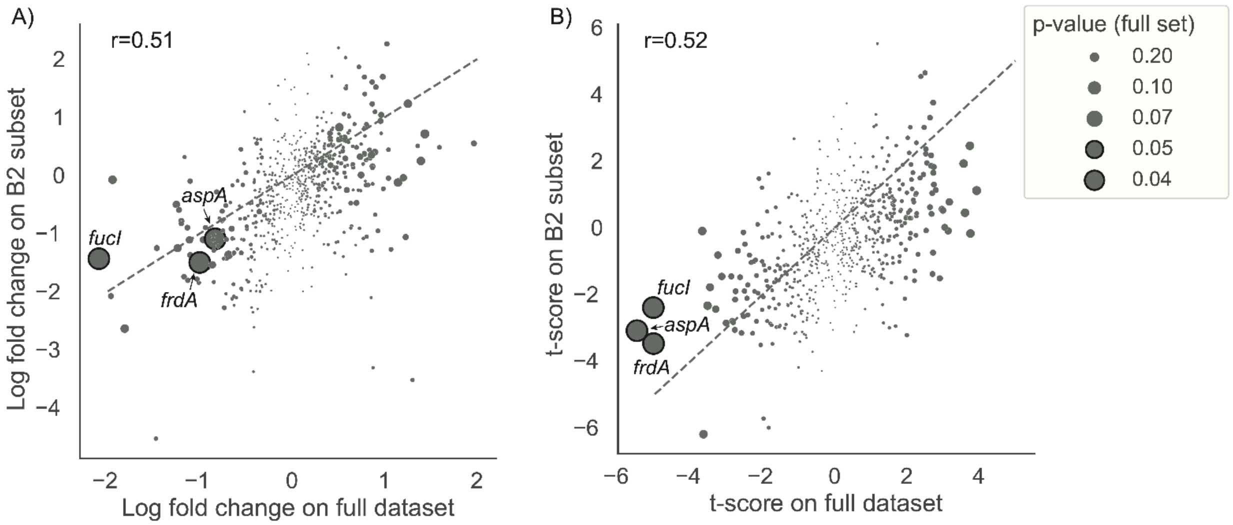


**Supplementary Figure 9. Results of phylogroup-controlled DE tests mirror results of full DE analysis**. DE models for 659 genes were re-fit to the 17 samples containing a single phylogroup B2 strain. Points represent DE coefficients from the full set of data (x-axis) and the B2 subset (y-axis). Points are sized by FDR-corrected p-value. Genes that were significant on the full set of data are labeled. A dashed line is drawn at y=x. a) estimated log fold changes (LFC) b) t-scores.

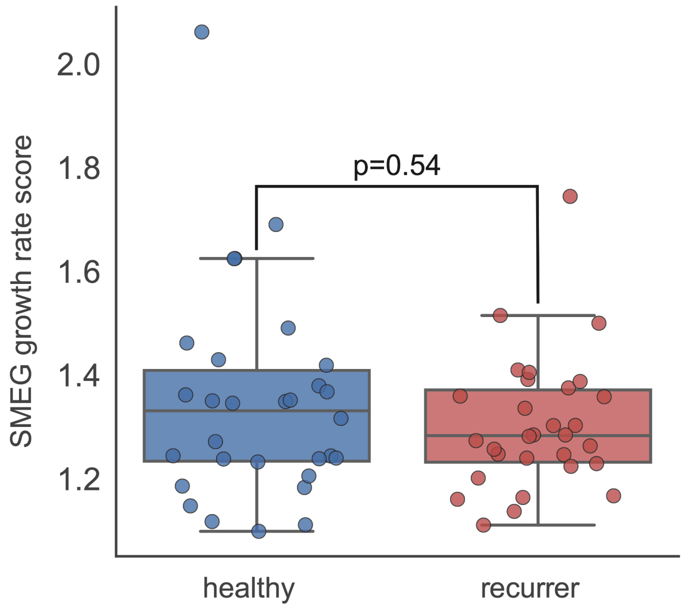


**Supplementary Figure 10. Estimated *E. coli* growth rate for samples from the healthy (blue) and recurrer (red) cohorts**. Growth rate was estimated as the difference in metagenomic coverage of the origin and terminus of replication, with SMEG (Methods).
